# CCL26 and CXCL12 Promote Release of Insulin-Sensitizing Adipose Tissue Macrophage sEVs from Subcutaneous Adipose Tissue in Obesity

**DOI:** 10.1101/2025.08.20.670151

**Authors:** Ishtiaq Jeelani, Flavia Franco da Cunha, Theresa V. Rohm, Chanond A. Nasamran, Jae-Su Moon, Letian Wang, Sruthika Prakash, Roi Issac, Cairo Murphy, Christopher K. Glass, Jerrold M. Olefsky, Yun Sok Lee

## Abstract

Unlike visceral adipose tissue (VAT), subcutaneous adipose tissue (SAT) can play a protective role against the development of insulin resistance and metabolic dysfunction in obesity. Here, we show that, in obesity, subcutaneous adipose tissue macrophages (ATMs) release small extracellular vesicles (sEVs) that can improve insulin sensitivity, opposite to the effect of visceral ATM sEVs. This functional difference was associated with an increase in the proportion of insulin-sensitizing, resident ATMs in SAT. *In vivo* and *in vitro* measurements of ATM growth and trafficking combined with single cell RNA sequencing (scRNA-seq) revealed that higher resident ATM survival and lower blood monocyte immigration along with decreased transition to pro-inflammatory ATMs collectively lead to the relative abundance of resident ATMs in SAT in obesity. These changes were mediated by CCL26 derived from subcutaneous adipocytes and adipocyte progenitors and CXCL12 secreted from resident ATMs. Our results elucidate previously unknown mechanisms for how SAT retains protective functions against metabolic dysfunction in obesity.

## Introduction

Obesity-induced chronic tissue inflammation is a major cause of insulin resistance and metabolic dysfunction ^1–4^. During the development of obesity, inflammation is initiated in adipose tissue and gradually propagates to other insulin target tissues, as well as pancreatic islets ^5,6^. In obese adipose tissue, macrophages account for the majority of the infiltrated immune cells and macrophages (especially, pro-inflammatory activated macrophages) are the major source of secreted factors that induce insulin resistance ^7–9^. Numerous studies have shown that depletion of macrophages or genetic manipulation to block pro-inflammatory macrophage activation improves insulin resistance and glucose tolerance in obese mice ^1,10–23^.

Notably, although obesity is typically associated with insulin resistance and metabolic dysfunction, some obese individuals remain insulin-sensitive and show a lower risk for developing type 2 diabetes mellitus (T2DM) and cardiovascular diseases ^24–29^. Although the molecular determinants for healthy vs unhealthy obesity remain to be elucidated, healthy obese subjects show preferential accumulation of body fat in lower body SAT (instead of VAT) depots ^30^, as exemplified by pear-shaped vs apple-shaped individuals ^31^. On the other hand, visceral adiposity is associated with inflammation and insulin resistance ^32^. Consistent with this concept, removal of VAT (omentectomy) decreases plasma glucose and insulin levels, whereas, removal of SAT (liposuction) does not ^33,34^. Moreover, the insulin-sensitizing effect of thiazolidinediones is associated with fat redistribution from VAT to SAT ^35,36^. Indeed, adipose tissue transplantation experiments in rodents revealed that SAT protects from metabolic abnormalities compared with VAT ^37,38^. Inflammation is lower in SAT with a smaller number of ATMs, compared with VAT ^39^, suggesting a mechanism in SAT that protects from the development of overt inflammation and metabolic dysfunction. Identifying this mechanism could provide a new opportunity for the development of novel therapeutic approaches.

In this study, we find that SAT ATMs from obese mice secrete small extracellular vesicles (sEVs) that can reverse insulin resistance; this is opposite to the effect of sEVs released from obese VAT ATMs which cause decreased insulin sensitivity. This functional difference was associated with a greater number of resident ATMs, which secrete sEVs that can improve insulin sensitivity. Mechanistic studies revealed that increased resident ATM survival, decreased blood monocyte immigration and transition to pro-inflammatory ATMs underly the favorable changes in SAT vs VAT ATMs. These effects were promoted by CCL26 and CXCL12 from SAT adipocytes and resident ATMs.

## Results

### SAT ATMs from obese mice secrete miRNAs that improve insulin sensitivity

In normal chow diet (NCD) healthy mice, VAT ATMs secrete sEVs that enhance insulin sensitivity, whereas, VAT ATMs in HFD/obese mice secrete sEVs that induce insulin resistance ^40^. To assess whether SAT and VAT ATMs secrete functionally different sEVs, we harvested sEVs released from ATMs isolated from SAT and VAT of NCD healthy and HFD obese mice (**Figure S1**), treated differentiated L6 myotubes with these ATM sEVs for 24h, and basal and insulin-stimulated glucose uptake was measured. Consistent with a previous report ^40^, healthy NCD VAT ATM sEVs enhanced insulin-stimulated glucose uptake (**Figure 1A**), whereas, HFD VAT ATM sEVs reduced insulin-stimulated glucose uptake without changing basal glucose uptake (**Figure 1B**). Similarly, NCD SAT ATM sEVs also increased insulin-stimulated glucose uptake (**Figure 1A**). However, unlike HFD VAT ATM sEVs, HFD SAT ATM sEVs did not induce insulin resistance (**Figure 1B**), and actually reversed the insulin resistance induced by palmitic acid (PA) treatment (**Figure 1C**, lane 16 vs 17).

**Figure 1.**
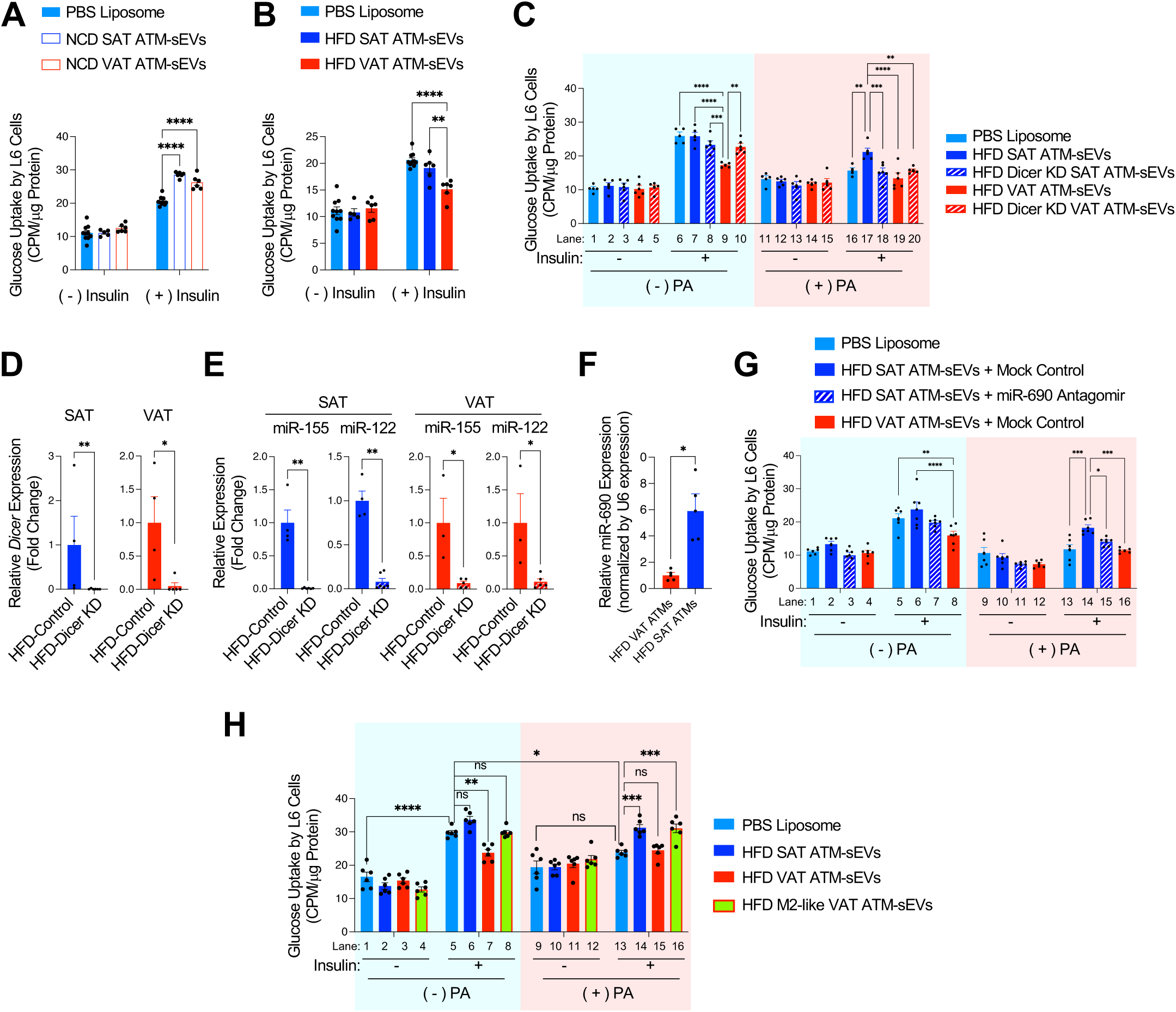
SAT ATMs secret sEVs that can improve insulin sensitivity in obesity. (**A**) Effect of sEVs derived from SAT and VAT ATMs of NCD mice on insulin sensitivity in normal L6 myotubes (n=6-9 wells/group). (**B**) Effect of sEVs derived from SAT and VAT ATMs of HFD mice on insulin sensitivity in normal L6 myotubes (n=5-10 wells/group). (**C**) Effect of DICER knockdown on the effect of HFD SAT and VAT ATM sEVs on regulating insulin sensitivity in normal and PA-treated, insulin resistant L6 myotubes (n=5 or 6 wells/group). (**D**) Relative *Dicer* expression ATMs (n = 4-6 wells/group). (**D**) Relative expression of miRNAs in control and *Dicer* KD ATMs (n= 3-6 wells/group). (**F**) Relative miR-690 expression in HFD SAT vs VAT ATMs (n=4 mice/group). (**G**) Effect of miR-690 antagomir treatment on insulin-sensitizing effect of HFD SAT ATM sEVs in normal and PA-treated, insulin resistant L6 myotubes (n=6 wells/group). (**H**) Effect of sEVs derived from SAT and VAT ATMs of HFD mice on insulin sensitivity in normal and PA-treated, insulin resistant L6 myotubes. Effects of sEVs released from CD11c^-^ CD206^+^ ATMs from visceral adipose tissue were compared in parallel (n=6 wells/group). ns, not significant. \**P* < 0.05, \*\**P* < 0.01, \*\*\**P* < 0.001, \*\*\*\**P* < 0.0001. All data are mean +/-SEM. Statistical analysis was performed by two-way ANOVA with Tukey’s multiple comparison tests.

The insulin-sensitizing, or desensitizing effects of ATMs are mediated, at least in part, by microRNAs (miRNAs), which are secreted in sEVs and taken up by insulin target cells such as adipocytes, hepatocytes, and myotubes ^41^. For example, miR-690 is enriched in sEVs from healthy NCD ATMs and mediates insulin-sensitizing effects of sEVs from these ATMs ^41,42^. DICER mediates the initiation of miRNA biogenesis ^43^ and DICER knockdown depletes miRNAs. To deplete microRNAs in ATM sEVs, we transfected HFD SAT and VAT ATMs with mock or *Dicer*-specific siRNAs and harvested sEVs secreted from these ATMs (**Figures 1D and 1E**). Consistent with previous reports ^41,42^, DICER knockdown abolished the effect of HFD VAT ATM sEVs to induce insulin resistance (lane 9 vs 10). Of interest, DICER knockdown also abolished the insulin-sensitizing effect of HFD SAT ATM sEVs (**Figure 1C**, lane 17 vs 18), suggesting that the insulin-sensitizing activities of HFD SAT ATM sEVs are mediated by miRNAs. This effect was associated with higher miR-690 expression in HFD SAT ATMs compared with HFD VAT ATMs (**Figure 1F**). Moreover, treatment with the miR-690 antagomir attenuated the effect of HFD SAT ATM sEVs to improve insulin resistance in PA-treated L6 myotubes (**Figure 1G**, lanes 14 vs 15). Together, these results suggest that the insulin-sensitizing effects of SAT ATMs are mediated, at least in part, by miR-690 contained in sEVs.

### Resident ATMs sEVs retain the insulin-sensitizing effect in obesity

NCD VAT is enriched in resident insulin-sensitizing ATMs, whereas HFD VAT is enriched in recruited ATMs, which can cause insulin resistance ^40^. However, HFD VAT still contains resident ATMs and it is not known whether obesity alters the insulin-sensitizing effect of resident ATMs. To address this question, we sorted CD206^+^ CD11c^-^ resident ATMs from total HFD VAT ATMs and compared the effect of sEVs released from these cells with sEVs from total HFD VAT ATMs on control and PA-treated insulin resistant L6 myotubes. As seen in **Figure 1H**, unlike total HFD VAT ATM sEVs, sEVs from CD206^+^ CD11c^-^ ATMs improved insulin sensitivity in PA-treated L6 cells (lanes 16 vs 13), similar to HFD SAT ATM sEVs (lanes 16 vs 14), suggesting that the beneficial effect of resident ATMs is maintained in obesity.

### Mechanism of how SAT contains relatively more insulin-sensitizing ATMs than VAT

The proportion of CD206^+^ CD11c^-^ ATMs is higher (47% vs 10%), whereas, the proportion of CD206^-^ CD11c^+^ recruited ATMs is lower (19% vs 33%) in SAT compared with VAT in HFD mice ^42^. Therefore, our results suggest that it is the differential proportion of resident ATMs in obese SAT vs VAT that determines the overall effect of total ATM sEVs on insulin sensitivity. Therefore, next, we focused on the mechanism of how insulin-sensitizing, resident ATMs selectively concentrate (in proportion) in SAT vs VAT in obesity. In obesity, increased ATM proliferation and increased migration of blood monocytes (which differentiate into recruited ATMs) to adipose tissue leads to ATM accumulation in VAT ^16,44,45^. When Ki67 positivity was used as a measure of ATM proliferation, HFD promoted proliferation of ATMs and Ly6C^+^ adipose tissue monocytes in both adipose tissue depots compared with NCD (**Figures 2A-2D** and **S2A**). These increases were greater than in liver (**Figures 2C-2D**). However, proliferation of CD11b^+^ F4/80^high^ ATMs or Ly6C^+^ F4/80^low^ monocytes was comparable in SAT vs VAT of HFD mice, although the number of ATMs was higher in VAT compared with SAT (**Figures 2A-2D**).

**Figure 2.**
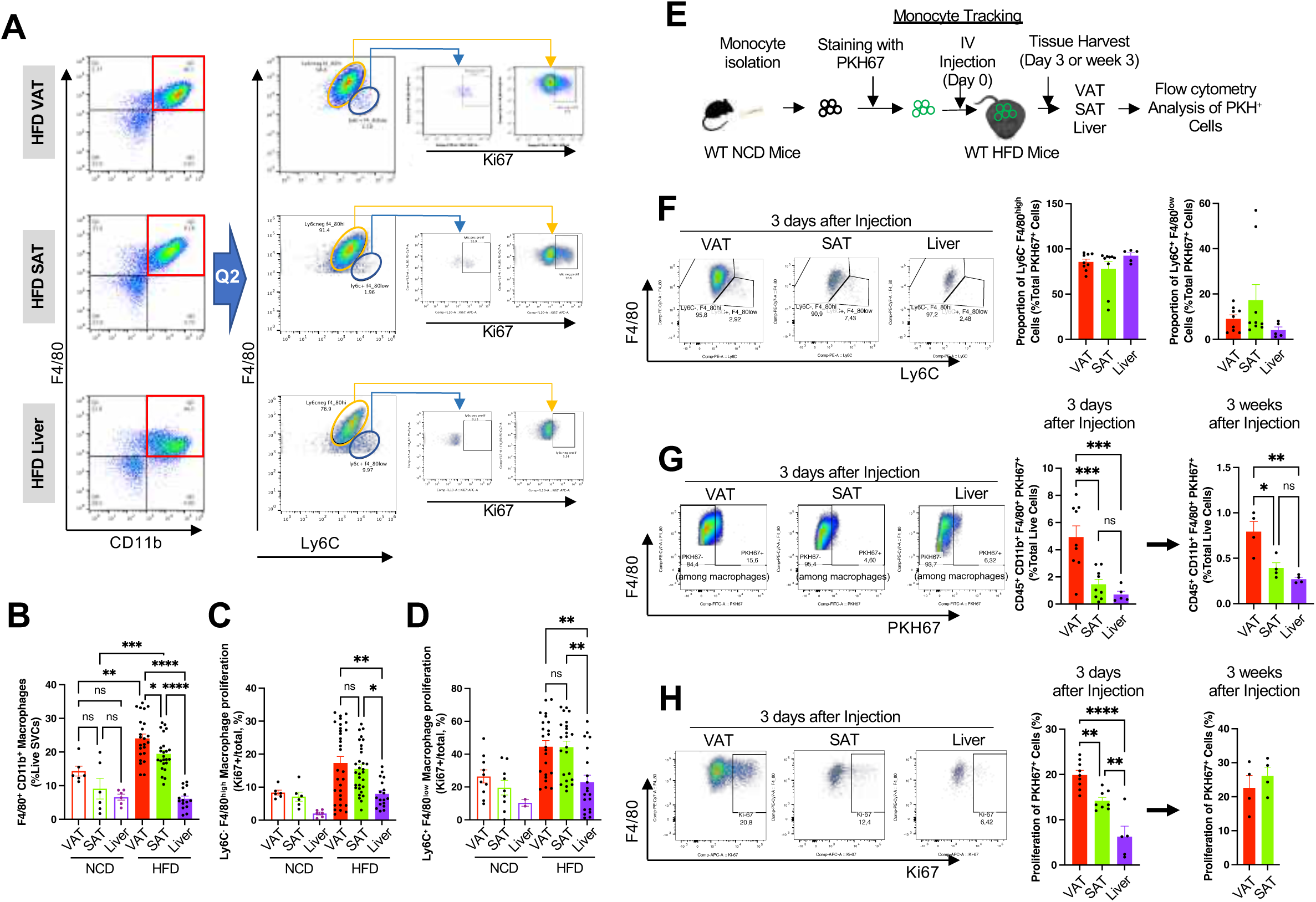
Blood monocyte immigration and newly recruited, but not all, ATM proliferation are lower in SAT compared with VAT in obesity. **(A-D)** Flow cytometry analysis of the proportion of total (**B**) and proliferating (Ki67^+^) (**C**) ATMs and Ly6C^+^ monocytes (**D**) in VAT and SAT of NCD/healthy and HFD/obese mice (n=2-30 mice/group). Representative blot images are shown in panel **A**. (**E-H**) Monocyte tracking experiments (n=4-9 mice/group). (**E**) Schematic representation of monocyte tracking experiments in mice. (**F**) Proportion of ATMs among PKH67-labeled cells three days after monocyte injection. (**G**) Proportion of PKH67-labeled, injected monocyte-derived ATMs in VAT and SAT of HFD/obese mice 3 days or 3 weeks after injection. (**H**) Proliferation of PKH67-labeled, recruited ATMs in VAT and SAT of HFD/obese mice 3 days or 3 weeks after injection. \**P* < 0.05, \*\**P* < 0.01, \*\*\**P* < 0.001, \*\*\*\**P* < 0.0001. All data are mean +/-SEM. Statistical analysis was performed by one-(G, H) or two-way (B-D) ANOVA with Tukey’s multiple comparison tests.

To assess monocyte immigration, we performed monocyte tracking experiments, as illustrated in **Figure 2E**. 3 days after PKH67-labeled monocyte injection, >80% of PKH67^+^ cells were Ly6C^-^ CD11b^+^ F4/80^high^ in both adipose tissue depots and liver (**Figure 2F**), indicating that the majority of injected monocytes that migrated to adipose tissue and liver had differentiated to macrophages over this time. Of interest, the proportion of injected monocyte-derived ATMs in the stromal vascular cells (SVCs) was higher in VAT compared with SAT and this change was sustained for up to 3 weeks (**Figure 2G**). Moreover, proliferation of newly migrated (3 day-old) monocyte-derived ATMs was higher in VAT compared with SAT (**Figure 2H**), although total ATM proliferation was comparable in HFD VAT and SAT (**Figure 2C**). This adipose depot-specific difference in recruited ATM proliferation was no longer present three weeks after PKH67-labeled monocyte injection (**Figure 2H**). Together, these results indicate that lower blood monocyte immigration and reduced proliferation of newly recruited ATMs contributes to the reduced ATM accumulation in SAT compared with VAT.

### Single cell RNA sequencing (scRNA-seq) analysis of ATMs

Our flow cytometry analyses showed that obesity can differentially affect proliferation of ATM or monocyte subtypes in the two adipose tissue depots, which can confer preferential resident ATMs concentration in SAT. To address this question at higher resolution, we performed scRNA-seq analyses in sorted live CD45^+^ CD11b^+^ myeloid cells in SAT and VAT from NCD and HFD mice. To understand whether obesity-induced changes in ATM transcriptomes are reversible, we also studied myeloid cells from adipose tissues of HFD mice after inducing improved insulin sensitivity by 3 weeks of a diet switch to NCD (weight loss) or 3 weeks of rosiglitazone (Rosi) treatment.

When all cells from all conditions were used, three different monocyte (Mon1∼3) and five different ATM (ATM1∼5) clusters with distinct transcriptomic signatures were identified (**Figures 3A-3C, S3A, and S3B**). A proliferating ATM/monocyte cluster was also identified. The ATM1 cluster was highly enriched in both SAT and VAT of healthy NCD mice (79∼84% of total ATMs) and declined on HFD, suggesting that ATM1 represents resident ATMs. Consistent with this, ATM1 was the only ATM cluster expressing a marker of tissue resident macrophages, *Timd4* (**Figure S3B**).

**Figure 3.**
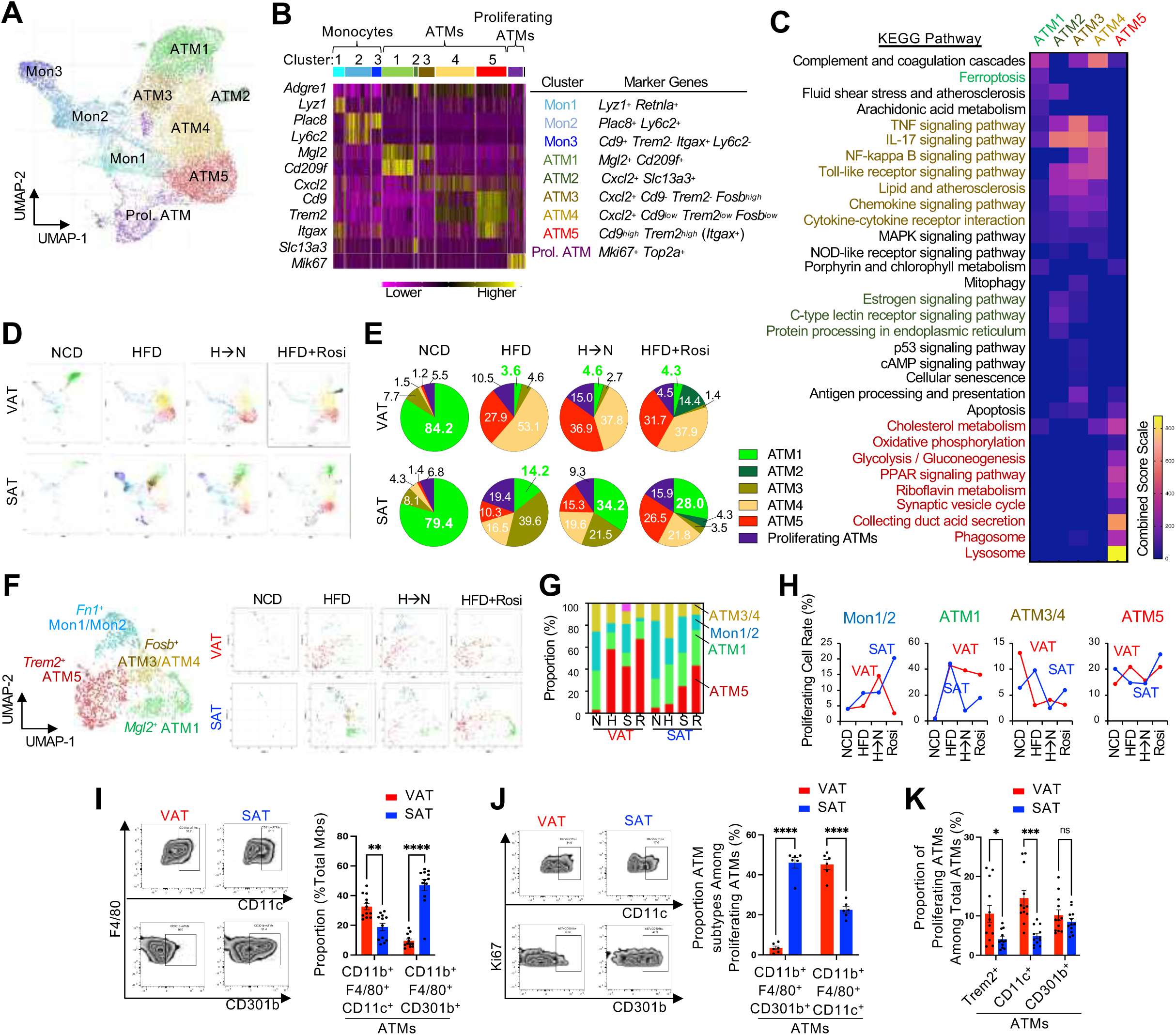
Subcutaneous adipose tissue provides microenvironment causing higher resident and lower recruited ATM proliferation compared with visceral adipose tissue in obesity. **(A-E)** Single cell RNA-seq analysis of ATMs in VAT and SAT of mice fed NCD, HFD, and HFD➔NCD switch diet, and HFD mice treated with rosiglitazone. (**A**) UMAP representation of six different ATM and three different monocyte clusters. (**B**) Heatmap showing cluster-specific marker gene expression in each of adipose tissue monocytes and macrophages. (**C**) Heatmap showing pathways enriched in each of five different ATM clusters. (**D**) UMAP representation of six different ATM and three different monocyte clusters in different diet and/or treatment conditions. (**E**) Proportion of each of six ATM clusters among total ATMs in different diet and/or treatment conditions. (**F**) UMAP representation of subclusters of the proliferating ATM cluster. Left, combined cell clusters from all conditions. Right, UMAP showing subclusters in each of different diet and/or treatment conditions. (**G**) Proportion of different ATM and monocyte clusters among all cell in the proliferating ATM cluster in each of different diet and/or treatment conditions. N, NCD; H, HFD; S, switch diet from HFD to NCD; R, Rosi treatment. (**H**) Proportion of proliferating cells in each of the ATM and monocyte clusters in each of different diet and/or treatment conditions.**(I-K)** Flow cytometry analysis of changes in the proportion of total and proliferating CD11c^+^ (recruited), Trem2^+^ (recruited), or CD301b^+^ (resident) ATMs in VAT and SAT of HFD mice. (**I**) Proportion of CD11c^+^ and CD301b^+^ ATMs among total ATMs (n=12 mice per group). (**J**) Proportion of CD11c^+^ (recruited) and CD301b^+^ (resident) ATMs among proliferating ATMs/monocytes (n=6 mice/group). (**K**) Proliferation of CD11c^+^ (recruited) and CD301b^+^ (resident) ATMs (n=12 mice/group). \**P* < 0.05, \*\**P* < 0.01, \*\*\**P* < 0.001, \*\*\*\**P* < 0.0001. All data are mean +/- SEM. Statistical analysis was performed by one-way ANOVA with Tukey’s multiple comparison tests.

Moreover, ATM1 was enriched in the expression of genes involved in anti-inflammatory activities, iron metabolism and complement and coagulation cascades (including *Mrc1* (encoding CD206), *Mgl2, Cd163,* and *Cd209f*) and expressed relatively lower levels of pro-inflammatory genes and did not express *Itgax* (encoding CD11c) (**Figures 3B and S3B**), similar to CD206^+^ CD11c^-^ ATMs that secreted insulin-sensitizing sEV. Therefore, we refer to ATM1s as healthy resident ATMs in the rest of this paper. Of note, on HFD, SAT showed relatively moderate decreases in the proportion of resident ATM1 (14%) compared with VAT (3.6%). Interestingly, the diet switch from HFD to NCD or treatment of HFD mice with Rosi increased the proportion of ATM1 in SAT (2.4-fold and 2-fold increases after diet switch or Rosi treatment, respectively), but not in VAT. This occurred concomitantly with SAT-specific decreases in the proportion of ATM3 (43% and 90%) after the diet switch and Rosi treatment, respectively) **(Figures 3D, 3E, and S3C**). The amount of ATM3, 4 and 5 clusters was negligible in NCD mice in both adipose depots, but was increased on HFD. These obesity-induced ATM clusters expressed *Cxcl2* and expression was significantly higher in VAT of HFD mice compared with VAT from NCD mice (**Figures S3B and S3D**). Pathway analysis using differentially expressed (DE) genes revealed that ATM3 and ATM4 were highly enriched in pro-inflammatory gene expression: Among these two proinflammatory ATM clusters, ATM3 showed a SAT-specific increase on HFD (from 8.1% of ATMs to 40%) and showed relatively higher expression of genes related to antigen presentation, cellular senescence, stress response, and apoptosis. On the other hand, ATM4 exhibited a stronger increase in VAT (35.3 fold) vs SAT (3.8 fold) on HFD, and showed higher levels of genes involved in cholesterol metabolism and complement and coagulation cascades. The ATM cluster also expressed both *Itgax* (often used to define proinflammatory, M1-like polarized ATMs) and *Mrc1* (**Figures 3C and S3B**), similar to the CD11c and CD206 double positive ATMs that are positively associated with HbA1c levels in humans ^46^. The ATM5 cluster did not display increased pro-inflammatory gene expression, but did show more abundant expression of genes involved in phagocytosis, lysosomal activity, lipid metabolism, and mitochondrial activity, including *Trem2* and *Cstd* (**Figures 3C and S3B**), similar to an ATM cluster previously named lipid-associated macrophages (LAMs) ^47^. Inflammatory gene expression was substantially lower in all three monocyte clusters (Mon1, 2, and 3) compared with ATM3 and ATM4 (**Figure S3E**). Together these results suggest that SAT maintains a higher number of healthy resident ATMs (ATM1) on HFD and the diet switch or Rosi treatment increased healthy resident ATMs in SAT but not in VAT. In contrast, VAT contained less resident ATMs and this did not reverse after the diet switch or Rosi treatment.

### Resident ATM proliferation is comparable in SAT and VAT in obesity

To assess whether the higher ratios between resident ATMs in SAT vs VAT were due to higher proliferation, we focused on the proliferating ATM cluster. The proliferating ATM cluster consisted of four major subclusters, showing expression of marker genes of ATM1 (*Mgl2*), ATM3/4 (*Fosb*), ATM5(*Trem2*), and Mon1/2(*Fn1*) (**Figure 3F**). On NCD, resident ATM1s accounted for the majority of proliferating ATMs in both SAT and VAT (**Figure 3G**). On HFD, the majority of proliferating ATMs became *Trem2*^high^ ATM5s in VAT and the ATM1 proportion was decreased, even after the diet switch or Rosi treatment. On the other hand, in SAT, although the proportions of ATM3/4 and ATM5 among total proliferating ATMs were increased on HFD, the increase was relatively modest compared with VAT (**Figure 3G**). Consequently, ATM1 comprised the majority of proliferating ATMs on HFD. However, the proportion of proliferating ATM1s among total ATM1s were comparable in HFD SAT and HFD VAT (**Figure 3H**), indicating that the larger number of resident ATM1s in SAT compared with VAT explains the greater number of proliferating ATM1s in SAT. Consistent with these results, flow cytometry analysis of ATMs from HFD mice showed that the proportion of CD301b^+^ (encoded by *Mgl2*, a marker of resident ATM1s) ATMs is higher and CD11c^+^ recruited ATMs is lower in SAT compared with VAT (**Figures 3I and S3F**). A majority of Ki67^+^ proliferating ATMs were CD301b^+^ resident ATMs in SAT, whereas, the majority of proliferating ATMs in VAT were recruited CD11c^+^ or Trem2^+^ ATMs (**Figures 3J and S3F**) ^45,47^. Moreover, the proportions of Ki67^+^ proliferating resident ATMs (CD301b^+^) among total resident ATMs were comparable in HFD SAT and VAT (**Figures 3E and 3F**), suggesting that proliferation of resident ATMs is not higher in SAT vs VAT.

### Mechanism of how SAT contains relatively more insulin-sensitizing ATMs than VAT (2): lower resident ATM death

To understand the mechanism by which HFD SAT maintains a relatively larger amount of resident ATMs (ATM1) compared with VAT in the absence of increased resident ATM proliferation, we analyzed DE genes between ATM1s in the two adipose tissue depots. Of interest, each of the five ATM clusters, including ATM1, showed many DE genes between SAT and VAT on each diet and/or treatment condition, although they still shared signature genes conforming to the same clusters (**Figure 4A**). For example, a group of genes enriched in VAT ATM1 compared with SAT ATM1 was associated with inflammatory pathways (e.g. TNF) in NCD mice (**Figure 4B**). On HFD, ATM1 in both adipose depots showed similar directional changes (**Figure 4C**), including increased expression of inflammatory cytokine and chemokine pathway genes. The increase was greater in SAT compared with VAT. Consequently, on HFD, SAT and VAT ATM1 showed similar levels of inflammatory gene expression and the number of DE genes between SAT and VAT ATM1 clusters decreased from 164 on NCD to 57 on HFD (**Figure 4A**). Pathway analysis of these 57 DE genes revealed that HFD VAT ATM1 expressed higher levels of genes involved in cellular stress responses, such as ferroptosis and mitophagy (**Figures 4D, 4E, and S4A**). This VAT-specific increase in stress response genes was specific to the ATM1 cluster (**Figures S4B-S4D**). These results raise the possibility that obesity induces preferential increases in cellular stress in resident ATMs (ATM1) in VAT compared with SAT, which might lead to increased resident ATM death. To assess whether obesity induces preferential increases in resident ATM death in VAT compared with SAT, we measured caspase-3/7 activity in ATMs sorted for the ATM1 marker (CD301b) from NCD and HFD mice. As seen in **Figure 4F**, HFD induced a higher increase in caspase-3/7 activity in CD301b^+^ ATMs in VAT, compared with SAT. To determine whether HFD induces the release of soluble factors that induce ATM death in VAT, we incubated BMDMs in HFD VAT or SAT-conditioned media (CM) for 24h and measured caspase-3/7 activity. As seen in **Figures 4G-4I**, HFD VAT, but not SAT, CM increased caspase-3/7 activity with increased lipid peroxide levels and the expression of genes involved in cell death. Treatment with the anti-oxidant ferrostatin that traps peroxy radicals did not block the pro-apoptotic effect of HFD VAT CM (**Figure 4G**).

**Figure 4.**
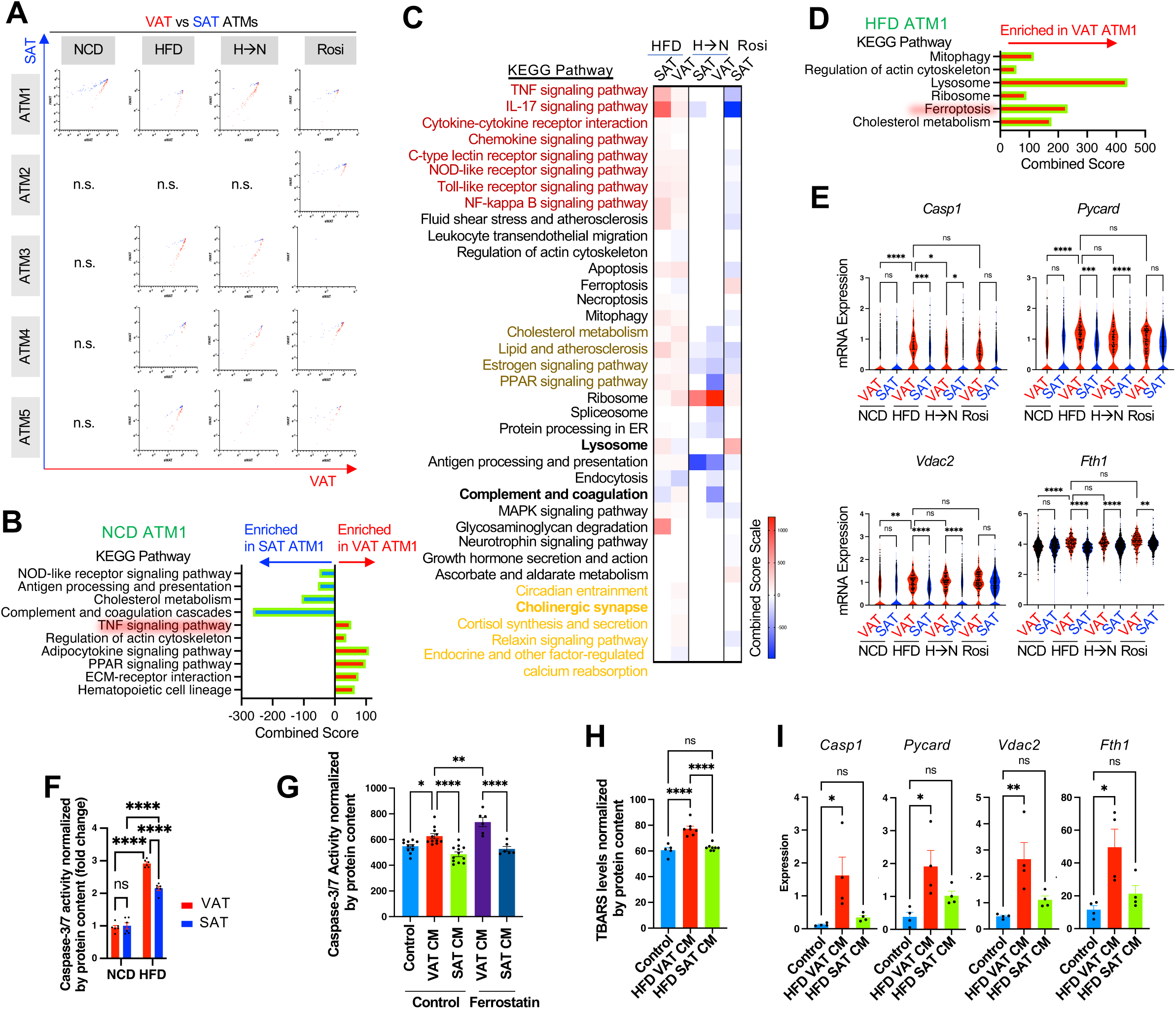
Subcutaneous adipose tissue provides microenvironment causing lower cell death of resident ATMs compared with visceral adipose tissue in obesity. **(A)** Scatter plots showing differential gene (DE) expression in each of five different ATM clusters residing in SAT and VAT in each of four different diet and/or treatment conditions. (**B**) Pathway analysis using DE genes in ATM1 from SAT and VAT of NCD mice. (**C**) Heatmap showing pathways that are up (red)- or down(blue)-regulated in ATM1 from SAT and VAT by HFD, the HFD➔NCD switch diet, or the treatment with Rosi. (**D**) Pathway analysis using DE genes in ATM1 from SAT and VAT of HFD mice. (**E**) mRNA expression of genes involved in cell death in ATM1 from SAT and VAT in each of different diet and/or treatment conditions. (**F**) Caspase-3/7 activity in sorted CD301b^+^ ATMs isolated from NCD and HFD mouse SAT and VAT (n=6 mice per group). (**G**) Effect of soluble factors released from obese VAT and SAT on macrophage apopotosis. BMDMs were incubated with HFD SAT- or HFD VAT-CM in the presence or absence of ferrostatin. 24h later, caspase-3/7 activities were measured (n=6-12 wells/group). (**H**) TBARS levels in BMDMs incubated in HFD VAT- or HFD SAT-CM (n=5-8 well/group). (**I**) Expression of genes involved in cell death in BMDMs incubated in HFD VAT- or SAT-CM (n=4 well/group). ns, not significant. \**P* < 0.05, \*\**P* < 0.01, \*\*\**P* < 0.001, \*\*\*\**P* < 0.0001. All data are mean +/- SEM. Statistical analysis was performed by 1- or 2-way ANOVA with Tukey’s multiple comparison tests.

### Transition of ATM1s to other clusters is not lower in SAT vs VAT

To assess the possibility that obesity induces enhanced transition of ATM1s to other ATM clusters in VAT compared with SAT, we conducted *in silico* lineage tracing using RNA Velocity and CytoTrace analyses. By calculating the ratio between unspliced vs spliced transcripts, these methods estimate the time derivative of the gene expression state and provide a snapshot of phenotypic switching. As seen in **Figures 4E-4K**, on NCD, this analysis indicated that ATM differentiation occurred through a linear trajectory from Mon1 and Mon2 to ATM1 in both VAT and that SAT and ATM1 was the end point of cellular transition. On HFD, transition of ATM1 toward ATM3 was seen in both VAT and SAT, and was somewhat greater in SAT. This suggests that the larger number of ATM1s in SAT compared with VAT was not due to decreased transition of ATM1s to other ATM clusters (such as ATM3).

### SAT-specific CCL26 expression increases insulin-sensitizing ATMs

Our data suggest that the larger amount of resident ATMs is an important determinant of the insulin-sensitizing effect of SAT (vs VAT) ATMs in obesity and this is due to higher resident ATM survival and lower blood monocyte chemotaxis in SAT compared with VAT (**Figures 2 and 4**).

Therefore, to identify factors causing these effects, we searched for soluble factors that are abundantly expressed in healthy SAT and VAT, but, are specifically decreased in obese VAT compared with obese SAT. Among these, we found that a repulsive chemokine ^48,49^, *Ccl26* is highly expressed in both SAT and VAT of NCD healthy mice (**Figures 5A and S5A**). On HFD, *Ccl26* expression was decreased in both SAT and VAT (**Figure 5A**). However, the decrease in VAT was greater. Consequently, *Ccl26* expression was substantially higher in HFD SAT compared with HFD VAT (**Figure S5B**). On the other hand, treatment with Rosi increased *Ccl26* expression in SAT of HFD/obese mice (**Figure 5A**). Of interest, this occurred selectively in SAT, but not VAT. The major source of *Ccl26* expression was adipocytes in NCD healthy mice. On HFD, adipocyte *Ccl26* expression was decreased and became comparable to that in SVCs (**Figure 5B**). Consistent with these results, *Ccl26* expression was higher in differentiated 3T3-L1 adipocytes compared with preadipocytes (**Figure 5C**). Rosi treatment did not affect *Ccl26* expression, consistent with the lack of effect of Rosi in NCD mice. Similar to HFD, increased metabolic stress induced by high glucose and PA levels decreased adipocyte *Ccl26* expression (**Figure 5D**), which was partially reversed by Rosi treatment (**Figure 5E**). Similarly, analysis of published human single nuclear (sn) RNA-seq data ^50^ revealed that CCL26 expression is greater in SAT compared with omental adipose tissue (**Figure S5C**). Unlike mice, in humans, CCL26 was more abundantly expressed in adipose stem and progenitor cells (ASPCs) compared with adipocytes (**Figure S5D**).

**Figure 5.**
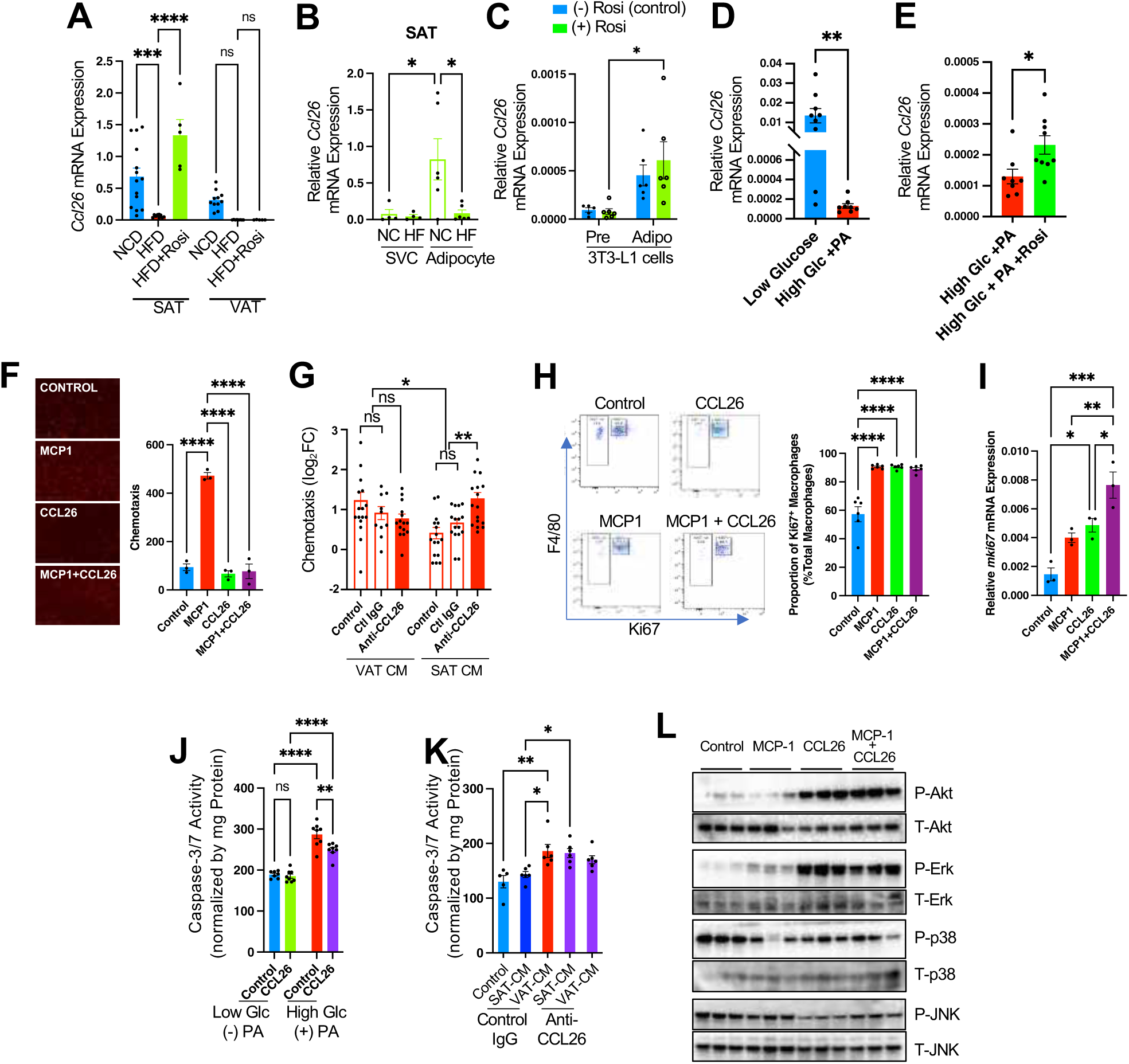
Subcutaneous adipocytes express CCL26 which blocks monocyte chemotaxis and increases macrophage proliferation and survival. **(A)** *Ccl26* mRNA expression in SAT and VAT of mice fed NCD or HFD, or HFD mice treated with rosiglitazone (Rosi) (n=14, 11, 4 mice/group). (**B**) *Ccl26* mRNA expression in stromal vascular cell (SVC) and adipocyte fraction of SAT from mice fed NCD (NC) or HFD (HF) (n=5 and 6 mice/group). (**C**) *Ccl26* mRNA expression in 3T3-L1 preadipocytes and differentiated adipocytes, treated with or without rosiglitazone (Rosi) (n=6 wells/group). (**D**) *Ccl26* mRNA expression in fully differentiated 3T3-L1 adipocytes incubated in low glucose or high glucose +PA medium for 24h (n=9 and 8 wells/group). (**E**) *Ccl26* mRNA expression in fully differentiated 3T3-L1 adipocytes incubated in high glucose +PA medium, +/- Rosi for 24h (n=9 and 8 wells/group). (**F**) Effect of CCL26 treatment on CCL2/MCP-1-induced monocyte (stained in red) chemotaxis. Representative images of migrated monocytes are shown on left (n=3 wells/group). (**G**) Effect of CCL26-neutralizing antibody treatment on chemoattraction by HFD VAT-CM and HFD VAT-CM (n=10 or 16 wells/group). (**H**) Effect of CCL26 on macrophage proliferation. 4 day-differentiated BMDMs were treated with or without 10 ng/ml MCP-1 and/or 100 ng/ml CCL26 for 24h. The proportion of Ki67^+^ proliferating macrophages were measured using flow cytometry (n=6 wells/group). (**I**) Effect of CCL26 on *mki67* expression in macrophages (n=3 wells/group). (**J**) Effect CCL26 on metabolic stress (high glucose+PA)-induced macrophage apoptosis (n=7 or 8 wells/group). (**K**) Effect CCL26 neutralization on macrophage apoptosis induced by factors released from obese VAT- or SAT-CM (n=5 or 6 wells/group). (**L**) Effect CCL26 treatment on cell signaling associated with cell survival and proliferation (Akt and MAPKs) in BMBMs (n=3 wells/group). \**P* < 0.05, \*\**P* < 0.01, \*\*\**P* < 0.001, \*\*\*\**P* < 0.0001. ns, not significant. All data are mean +/- SEM. Statistical analysis was performed by the student t-tests (D,E) or 1 or 2-way ANOVA (A-C, F-K) with Tukey’s multiple comparison tests.

CCL26 binds to various chemokine receptors, such as CCR1, CCR2, and CCR5 and antagonizes monocyte chemotaxis ^48,49^. To assess whether CCL26 can suppress immigration of blood monocytes to adipose tissue, we performed monocyte chemotaxis assays in Transwell plates. Consistent with previous reports ^48,49^, CCL26 inhibited macrophage chemotaxis to MCP-1 (**Figure 5F**), which is a major chemokine inducing monocyte migration to adipose tissue in obesity ^4^. To assess whether CCL26 affects changes in the immigration of blood monocytes to SAT vs VAT in obesity, we measured monocyte chemotaxis to HFD SAT-CM and HFD VAT-CM in the presence or absence of anti-CCL26 neutralizing antibodies in Transwell plates. As seen in **Figure 5G**, treatment with CCL26 neutralizing antibodies increased monocyte chemotaxis to SAT CM to comparable levels induced by VAT CM.

MCP-1 also mediates obesity-induced resident ATM proliferation ^44^, as well as monocyte chemotaxis. Therefore, we assessed whether CCL26 affects proliferation and/or survival of macrophages. Since the number of resident ATMs is limited in mice, to obtain sufficient amounts of cells, we used BMDMs in these studies. Interestingly, treatment with CCL26 increased BMDM proliferation and the expression of a marker gene of proliferating cells, *mki67,* independent of MCP-1 (**Figures 5H, 5I, and S5E**). In addition, CCL26 treatment also inhibited metabolic stress-induced apoptotic caspase-3/7 activity in BMDMs (**Figure 5J**), while CCL26 neutralization enhanced the effect of HFD SAT CM to increase macrophage apoptosis (**Figure 5K**). These effects were associated with increases in the signaling pathways involved in cell growth and survival, such as Akt and Erk (**Figure 5L**). Taken together, these results suggest that SAT adipocytes release CCL26, which enhances ATM survival and proliferation and blocks blood monocyte immigration.

### SAT-specific CXCL12 expression promotes the secretion of insulin-sensitizing sEVs

We also assessed whether the larger resident ATM population in obese SAT provides a microenvironment affecting the SAT-specific changes to increase the number of insulin-sensitizing ATMs. Since both SAT and VAT are enriched in resident ATMs (ATM1) in NCD healthy mice, we isolated ATMs from both depots in NCD/healthy mice and harvested ATM-CM. As seen in **Figure S6A**, monocyte chemotaxis assays using Transwell plates revealed that healthy ATM-CM did not block MCP-1-stimulated monocyte chemotaxis. To assess whether resident ATMs regulate the differentiation and/or polarization states of recruited ATMs, we co-cultured partially differentiated (4 days) BMDMs (lower chamber) with resident ATMs (upper chamber) in Transwell plates and measured marker genes of monocytes and different ATM clusters in the BMDMs. As seen in **Figure 6A**, Transwell coculture of BMDMs with resident ATMs decreased the expression of macrophage markers in BMDMs such as *Trem2* and *Cd9* (enriched in ATM5) and increased the expression of a monocyte marker, *Ly6c2* (enriched in Mon2). Consistent with this, incubation of BMDMs with resident ATM CM induced decreased expression of *Adgre1* (encoding F4/80), *Trem2*, and *Itgax* (encoding CD11c), and increased the expression of *Ly6c2* and *Fosb* (enriched in ATM3) (**Figure 6B**), suggesting that resident ATMs can affect the differentiation and polarization states of newly recruited ATMs by releasing soluble factors.

**Figure 6.**
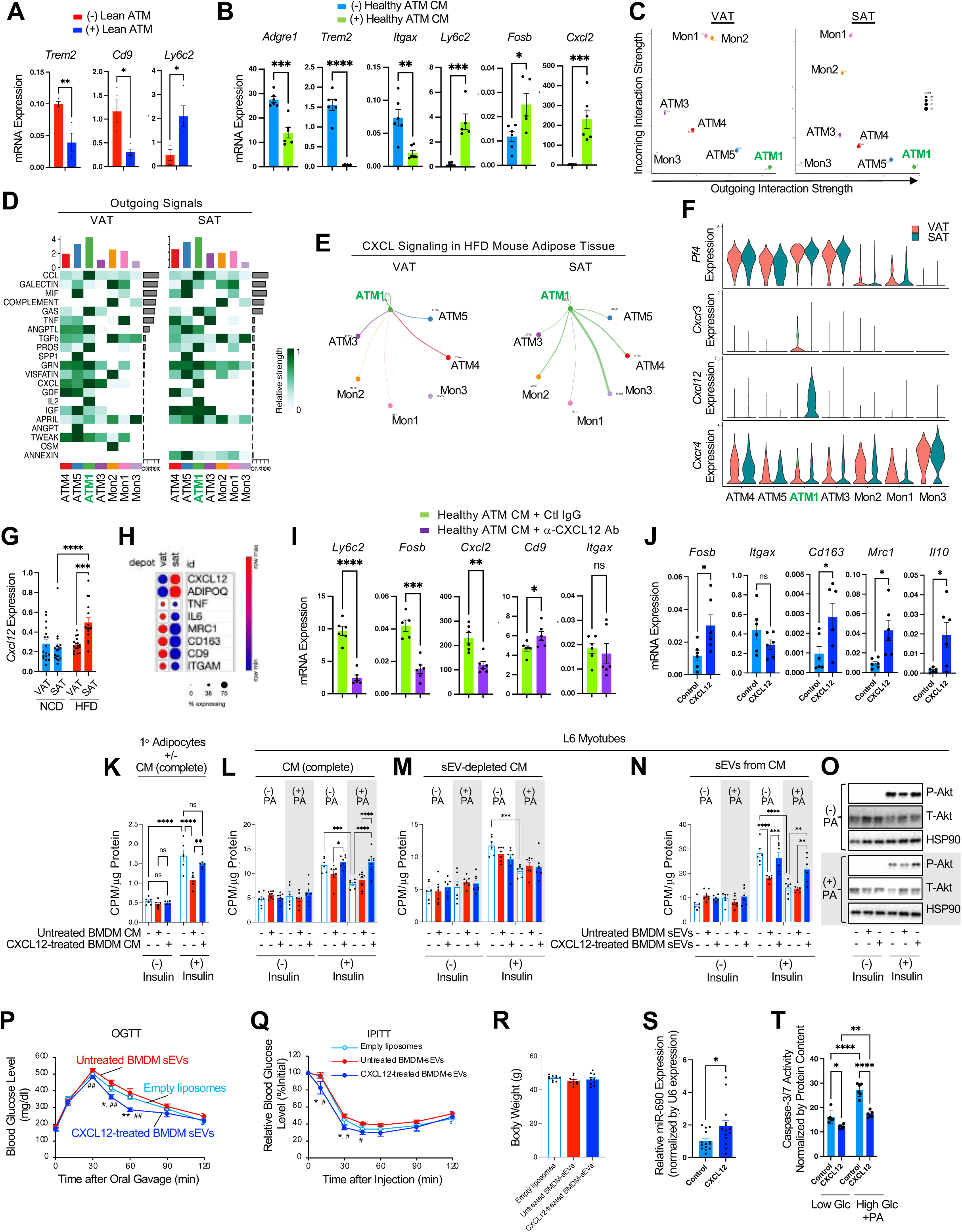
Subcutaneous resident ATMs express CXCL12 that can increase ATM survival and prevent from other ATMs from releasing soluble factors that induce insulin resistance. (**A**) Macrophage and monocyte marker gene expression in 4 day-differentiated BMDMs cocultured with healthy ATMs. (**B**) Macrophage and monocyte marker gene expression in 4 day-differentiated BMDMs incubated in healthy ATM-CM. (**C-F**) CellChat Analysis of interaction between different adipose tissue monocyte and ATM clusters in HFD mice. (**C**) Incoming and outgoing interaction strength of each ATM cluster in VAT (left) and SAT (right) of HFD mice. (**D**) Heatmap showing strength of outgoing signals from each of individual adipose tissue monocyte and ATM clusters. (**E**) Circle plot showing interaction between individual adipose tissue monocyte and ATM clusters through CXCL signaling. (**F**) Violin plot showing the expression of genes involved in CXCL signaling in each of adipose tissue monocyte and ATM clusters from HFD mouse SAT and VAT. (**G**) mRNA expression of *Cxcl12* in SAT and VAT of mice fed NCD or HFD (n=16 mice/group). (**H**) A dot plot showing relative mRNA expression of *Cxcl12* in human SAT and VAT (n=5 and 6 individuals/group)^53^. (**I**) Macrophage and monocyte marker gene expression in 4 day-differentiated BMDMs incubated with healthy ATM-CM, +/- anti-CXCL12 antibodies for 24h (n=6 wells/group). (**J**) Macrophage and monocyte marker gene expression in 4 day-differentiated BMDMs treated with or without CXCL12 for 24h (n=6 wells/group). (**K-O**) Insulin-stimulated glucose uptake (J-M; n=6 wells/group) and Akt phosphorylation (O) by primary adipocytes from HFD/obese WT mice (**K**) or L6 myotubes (**L**-**O**). Before measurements, cells were incubated in CXCL12-treated or untreated BMDM-CM (**K-L**) or CXCL12-treated or untreated BMDM-derived sEVs (**N-O**) for 24h. To test the effect of sEVs in the mediation of the effect of BMDM-CM, we used both complete (**K, L**) and sEV-depleted BMDM-CM (**M**). (**P-R**) Effect of CXCL12-treated BMDM sEVs on glucose (P) and insulin (Q) tolerance and body weight (R) in HFD mice (n=10 mice/group). (**S**) Relative miR-690 expression in BMDMs treated with or without CXCL12 for 24h (n=14 wells/group from three different batches of experimental samples). (**T**) Caspase-3/7 activity in BMDMs incubated in low or high glucose+PA media in the presence or absence of 10 ng/ml CXCL12 for 24h (n=6 wells/group). \**P* < 0.05, \*\**P* < 0.01, \*\*\**P* < 0.001, \*\*\*\**P* < 0.0001 vs Empty liposomes or as indicated. ^#^*P* < 0.05, ^##^*P* < 0.01 vs untreated BMDM sEVs. All data are mean +/- SEM. Statistical analysis was performed by the Student t-tests (A, B, H, I, R) or 1- or 2-way ANOVA with Tukey’s multiple comparison tests (G, J-M, S).

To explore the underlying mechanism, we conducted CellChat analysis in HFD mice. CellChat analyzes the expression of ligands and their specific receptors and coreceptors to quantitatively infer intercellular communication networks from scRNA-seq data ^51^. As seen in **Figure 6C**, ATM1 (resident ATMs) showed the strongest outgoing interaction strength, compared with other ATM and monocyte clusters. In SAT, the interaction strength of ATM1 with other ATMs was strong, although with fewer modules, compared with VAT (**Figures S6B and S6C**). These modules included the CXCL pathway (**Figure 6D**). Among CXCL pathway components, *Cxcl12* expression was highly enriched in SAT ATM1 compared with VAT ATM1 so that CellChat analysis showed interactions between ATM1 and other ATM clusters and monocytes in SAT, but not in VAT (**Figures 6E, 6F, and S6D**). Moreover, overall *Cxcl12* expression was higher in SAT compared with VAT in HFD mice (**Figure 6G**). Consistent with these results, snRNA-seq data ^50^ analysis revealed that *Cxcl12* was abundantly expressed in resident ATMs from obese SAT, but not VAT, of HFD male mice (**Figures S6E-S6H**). Among two CXCL12 receptors (CXCR4 and ACKR3) ^52^, adipocytes and adipocyte progenitors expressed *Ackr3* (but not *Cxcr4*), whereas, most immune cells including ATMs expressed *Cxcr4,* but not *Ackr3* (**Figures S6E-S6G**), suggesting that CXCL12 transmits intracellular signals through different receptor systems in adipocytes and ATMs. Similarly, analysis of published snRNA-seq data of human adipose tissue ^50,53^ revealed that *CXCL12* expression was higher in SAT compared with VAT (**Figures 6H and S6I**) and *CXCL12* was more abundantly expressed in *MRC1(CD206)*^+^ *LYVE1*^+^ *ITGAX(CD11c)*^-^ *TREM2*^-^ resident ATMs (hMac1) compared with *MRC1*^-^ *LYVE1*^-^ *ITGAX*^+^ *TREM2*^+^ recruited ATMs (hMac2) (**Figures S6J-S6K**).

To assess whether CXCL12 regulates ATM differentiation, we pre-incubated NCD healthy mouse ATM (enriched with ATM1) CM with or without CXCL12 neutralizing antibody and treated 4 day differentiated BMDMs. As seen in **Figure 6I**, CXCL12 neutralization reversed the effect of healthy ATM CM to increase *Ly6c2*, *Cxcl2*, and *Fosb* expression without changing *Itgax* levels. Consistent with this, CXCL12 treatment increased the expression of *Fosb, Trem2,* and anti-inflammatory macrophage marker genes such as *Mrc1, Il10*, and *Cd163* without changing *Itgax* expression (**Figure 6J**). These results suggest that resident SAT ATMs can change the polarization states of other ATMs, at least in part, by releasing CXCL12.

To test whether CXCL12 modulates the effect of ATMs on insulin sensitivity, we treated BMDMs with or without recombinant CXCL12 and harvested control and CXCL12-treated BMDM-CM (free of recombinant CXCL12), as illustrated in **Figure S6L**. Insulin resistant primary mouse adipocytes were isolated from HFD/obese WT mice and incubated with the CM for 24h, before measuring glucose uptake. Unlike control BMDM-CM, CXCL12-treated BMDM-CM did not impair insulin-stimulated glucose uptake (**Figure 6K**). Direct treatment of obese adipocytes with recombinant CXCL12 did not change insulin sensitivity (**Figure S6M**). Similarly, L6 myotubes incubated with CXCL12-treated BMDM-CM displayed improved insulin-stimulated glucose uptake compared with L6 myotubes incubated with untreated BMDM-CM (**Figures 6L and S6N**). Of interest, in the absence of PA, CXCL12-treated BMDM-CM did not enhance glucose uptake compared with untreated condition. However, in PA-treated insulin resistant L6 myotubes, CXCL12-treated BMDM-CM reversed PA-induced insulin resistance (**Figure 6L**). To assess whether CXCL12 treatment increased macrophage secretion of sEVs that enhance insulin sensitivity, we purified sEVs from CXCL12-treated and untreated BMDM-CM (**Figure S6O**) and also prepared CXCL12-treated and untreated BMDM-CM that were depleted of sEVs. Of interest, sEV depletion of CXCL12-treated BMDM-CM erased its effect to increase insulin sensitivity (**Figure 6M**). On the other hand, treatment with sEVs secreted from CXCL12-treated BMDMs improved insulin-stimulated glucose uptake, similar to the effect of complete CM (**Figure 6N**), with increased Akt phosphorylation (**Figure 6O**).

To test this concept in vivo, we treated HFD/obese mice with untreated control BMDM- or CXCL12-treated BMDM-sEVs and measured insulin and glucose tolerance. As seen in **Figures 6P-6R**, CXCL12-treated BMDM-sEVs improved insulin and glucose tolerance without changing body weight compared with empty liposomes or untreated BMDM-sEVs. These effects of CXCL12-treated BMDM sEVs was associated with a larger amount of miR-690 compared with untreated BMDM sEVs (**Figures 6S and S6P**). Lastly, we tested whether CXCL12 affects macrophage survival. As seen in **Figure 6T**, CXCL12 treatment reduced apoptotic caspase-3/7 activity induced by chronic incubation in high glucose + PA medium. Together, these results suggest that healthy ATMs secrete CXCL12, which can increase ATM survival and enhance secretion of insulin-sensitizing sEVs from other ATMs.

## Discussion

SAT plays a protective role against the development of insulin resistance and metabolic dysfunction in obesity, whereas, VAT has the opposite effect. However, the underlying mechanisms are not clearly understood. In this paper, we demonstrate that, compared with VAT, SAT contained a larger resident ATM population which secretes beneficial sEVs that promote insulin-sensitivity. In obese mice, resident ATM survival was increased in SAT compared with VAT. This was accompanied by lower blood monocyte immigration, decreased transition to pro-inflammatory ATMs, and a lower rate of proliferation of newly recruited ATMs. As a SAT-specific factor mediating these effects, we found CCL26 secreted from subcutaneous adipocytes (mice) and adipocyte progenitors (humans) promoted macrophage survival and blocked blood monocyte immigration. Moreover, we found healthy resident ATMs (ATM1s) in SAT secreted CXCL12 which enhanced release of insulin-sensitizing sEVs. Together, our results suggest that higher resident ATM survival and lower blood monocyte immigration induced by CCL26 and CXCL12 concentrate insulin-sensitizing, resident ATMs in obese SAT vs VAT and retain the protective effect against the development of insulin resistance in obesity.

Previously, it was suggested that recruited ATMs secrete factors that induce insulin resistance and, therefore, the increase in recruited ATMs are responsible for the development of insulin resistance in obesity. Therefore, previous studies about ATMs were highly focused on the mechanism for how obesity increases the number and function of proinflammatory, insulin resistance-inducing ATMs. However, this concept was challenged by later observations that improved insulin sensitivity after insulin-sensitizing regimens (such as weight loss or thiazolidinedione treatment) precedes regression of recruited ATMs. Moreover, clinical studies targeted to proximal inflammatory pathways showed disappointing results. In this study, we found that, although the proportion of resident ATMs was decreased in VAT in obesity, they retained the insulin-sensitizing effects. Of interest, compared with VAT, SAT maintained a four-times larger resident insulin-sensitizing ATM population and SAT ATM-derived sEVs improved PA-induced insulin resistance. Of note, the proportion of SAT resident ATMs, but none of recruited ATM clusters (e.g. ATM4 and ATM5) in both SAT and VAT, was precisely associated with changes in metabolic profile in mice after HFD, the switch diet from HFD to NCD, or Rosi treatment. These results suggest the possibility that the obesity-induced decrease in insulin-sensitizing resident ATMs, as well as the increase in insulin resistance-inducing recruited ATMs, contributes to the development of insulin resistance.

Several factors exhibiting differential expression between obese SAT and VAT were proposed to mediate the preferential increase in inflammation in obese VAT compared to SAT. However, SAT-specific factors that increase insulin-sensitizing, resident ATMs are not known. In this study, we discovered that CCL26 and CXCL12 act as SAT-specific factors promoting the relative abundance of insulin-sensitizing, resident ATMs in obesity. CCL26 is a member of the eotaxin subfamily of CC chemokines ^54–56^. Unlike other isoforms, CCL26 is mainly expressed in non-hematopoietic cells and is induced by anti-inflammatory Th2 cytokines ^57–59^. Moreover, CCL26 binds to various chemokine receptors, such as CCR1, CCR2, and CCR5 ^60^ and inhibits monocyte chemotaxis ^48,49^. In this study, we found that CCL26 neutralization increased monocyte chemotaxis to HFD SAT CM so it was now comparable to HFD VAT CM-induced chemotaxis. Moreover, CCL26 treatment directly decreased macrophage apoptosis induced by chronic high glucose and PA levels. Based on these findings, we suggest that CCL26 contributes to the relative abundance of resident ATMs in SAT vs VAT in obesity by inhibiting monocyte chemotaxis and promoting resident ATM growth.

CXCL12 is one of the most studied chemokines, mediating various physiologic functions, including macrophage trafficking, survival, differentiation and phagocytosis ^61–65^. Similar to CCL26, CXCL12 can also mediate chemorepulsion of T cells ^66^. In VAT and brown adipose tissue (BAT), *Cxcl12* is mainly expressed in pericytes and smooth muscle cells and increases UCP1 expression and energy expenditure. Deletion of *Cxcr4* in aP2- or UCP1-expressing cells (adipocytes) exaggerates HFD-induced body weight gain, glucose intolerance, and insulin resistance ^67,68^. Consistent with this, deletion of *Cxcl12* in *Acta2*-expressing pericytes and smooth muscle cells reduces energy expenditure and enhances weight gain on HFD in mice ^69^. In this study, we found that *Cxcl12* was preferentially expressed in SAT (but not VAT) resident ATMs of obese mice and controls differentiation of recruited ATMs. Moreover, CXCL12 treatment promoted the secretion of insulin-sensitizing sEVs in BMDMs. sEVs from CXCL12-treated BMDMs increased insulin-stimulated glucose uptake and Akt phosphorylation in L6 myotubes and improved glucose and insulin tolerance in HFD/obese mice. Therefore, along with the anti-obesogenic effect (increased thermogenesis), it is likely that CXCL12 can enhance insulin sensitivity by inducing macrophage differentiation to insulin-sensitizing phenotypes.

In contrast, it was also shown that deletion of *Cxcl12* in adiponectin-expressing mature adipocytes decreases body weight and adipose tissue mass, with a corresponding improvement in glucose and insulin tolerance ^70^. A possible explanation for these contradicting results may include that, depending on the source of production, CXCL12 can exert different metabolic effects.

CXCL12 cleavage at its N-terminus by exopeptidases (including DDP4) disables its effect to activate CXCR4 without altering its binding to CXCR4, making it a competitive CXCR4 antagonist^71^. Moreover, N-terminus cleavage of CXCL12 does not affect its ability to activate ACKR3. *Dpp4* is expressed in white adipocytes and ASCs, but not resident ATMs, and is increased in obese VAT compared with lean VAT or obese SAT ^72,73^. Therefore, it is possible that differential post-translational modification of CXCL12 in different cellular origins may affect different metabolic effects through differential activation of CXCR4 and ACKR3. Of interest, our results indicate that white adipocytes and adipocyte progenitors express *Ackr3*, but not *Cxcr4,* whereas ATMs expressed *Cxcr4* but not *Ackr3*, suggesting that white adipocytes and ATMs receive CXCL12 signal through different receptors. Future studies are required to address this question.

Comparative analysis of scRNA-seq data revealed that SAT and VAT displayed similar ATM profiles on NCD. However, ATMs in the two adipose tissue depots showed markedly different responses to HFD/obesity, to the diet switch from HFD to NCD and to Rosi treatment. On HFD, the proportion of healthy resident ATMs (ATM1) was much higher in SAT compared with VAT. In addition, HFD induced differential changes in the proportion of all three obesity-induced ATM clusters (i.e. ATM3, 4 and 5) in SAT vs VAT. Among the three obesity-induced ATM clusters, ATM3 represented the major pro-inflammatory ATMs in SAT, whereas, ATM4 represented the major pro-inflammatory ATMs in VAT. ATM4 and ATM5 cells were enriched in *Trem2* and *Itgax* expression, consistent with previous reports that obesity induces Trem2^+^ and CD11c^+^ ATMs in VAT ^47^ ^9,47^. On the other hand, SAT-specific ATM3 cells did not express *Itgax* or *Trem2*, suggesting this is a previously unidentified ATM population among obesity-induced pro-inflammatory ATMs. *In silico* lineage tracing indicated that ATM3s were derived, in part, from healthy resident ATMs (ATM1s). Since ATM1 death was lower in SAT compared with VAT, it is possible that increased ATM1 survival in SAT led to the transition of ATM1s to pro-inflammatory ATM3s in SAT. Indeed, after Rosi treatment, or the diet switch from HFD to NCD, the proportion of ATM3s was decreased along with increased ATM1s in SAT. This was accompanied by reduced transition of ATM1s to ATM3s. These results show there are different developmental lineages of the major pro-inflammatory ATM subtypes in SAT (ATM3) and VAT (ATM4). Furthermore, the lower SAT ATM1 death in obesity conferred faster recovery of healthy resident ATMs after the diet switch or Rosi treatment in SAT compared with VAT. In addition to the differences in the proportion of ATM subtypes in SAT vs VAT, each of the ATM clusters showed depot-specific differential gene expression. In particular, DE genes in both ATM3 and ATM4 in SAT vs VAT largely included genes associated with macrophage activity and metabolism. Future studies are required to understand whether these gene expression changes lead to functional differences in each ATM cluster in SAT vs VAT.

Since female mice develop relatively mild obesity and insulin resistance on HFD, our studies were conducted in male mice. Of interest, unlike HFD male mice, in HFD female mice, *Cxcl12* was expressed in both SAT and VAT resident ATMs. Therefore, it would be of interest in the future to determine the extent to which the depot specific changes in SAT and VAT described in this paper are expressed in female mice. The tissue cues or signals, causing adipocytes to secrete CCL26 or ATMs to secrete CXCL12, are not clear and remain to be defined with further work.

In summary, here we demonstrate a previously unknown mechanism by which SAT is protected from the development of inflammation in obesity. Changes in resident ATMs in obesity are an understudied area and our data suggest that obesity induces differential changes in growth and function of resident ATMs in SAT vs VAT. For example, we found sEVs from HFD SAT ATMs caused beneficial effects to improve insulin sensitivity in obesity, compared with HFD VAT ATM sEVs. Moreover, we identified ATM3 as a unique SAT-specific pro-inflammatory ATM cluster which is partly derived from resident ATMs. Our results add to our understanding of obesity-induced inflammation and may point the way towards the development of novel insulin-sensitizing approaches.

## Resource Availability

### Lead contact

Requests for further information and resources should be directed to the lead contact, Yun Sok Lee, who will handle and respond to them accordingly.

### Materials availability

This study did not generate new unique reagents.

### Data and code availability

The RNA-seq data have been deposited at GEO (GSE267767)

All other data reported in this paper will be shared by the lead contact upon request. This paper does not report original code.

## Acknowledgements

We thank Laiba Mohyuddin for helping BMDM preparation and realtime RT-PCR. This study was supported by the National Institute of Diabetes and Digestive and Kidney Diseases of the National Institutes of Health (NIH) (DK124298, DK120515, DK063491), the National Heart, Lung, and Blood Institute (HL142214), a UCSD Health Sciences Research Grant (RG084153), and a grant from Janssen Pharmaceuticals, Inc. This publication includes data processed at IGM Genomics Center and Center for Computational Biology and Bioinformatics (UL1TR001442 of CTSA). The content is solely the responsibility of the authors and does not necessarily represent the official views of the NIH. T.V.R. was supported by Larry Hillblom postdoc fellowship (2023-D-012-FEL) and Swiss National Science Foundation Postdoc Mobility grant (P2BSP3_200177).

## Author Contributions

I.J., F.F.C, T.V.R designed experiments and collected, analyzed, and interpreted data. L.W., S.P. prepared BMDMs and supported chemotaxis, realtime RT-PCR, CM and sEV preparation, and/or flow cytometry experiments. C.M. conducted glucose uptake experiments in L6 cells. C.A.N. supported scRNA-seq data analysis. C.K.G. contributed to the analysis and interpretation of data. J.M.O supervised the project, interpreted data, and wrote the manuscript. Y.S.L. conceived and supervised the project, designed experiments, collected, analyzed, and interpreted data, and wrote the manuscript.

## STAR Methods

### Animals and treatments

Male wild type C57BL/6J were purchased from Jackson Laboratory and housed in colony cages maintained at room temperature (22 °C) on a 12 h light/12 h dark cycle. 8 week-old male mice were fed either a lard-based 60% HFD (Cat# D12492, Research Diets) or a chow control diet for 16-20 weeks ad libitum. Since female mice only develop modest obesity and insulin resistance on a HFD, male mice were used in all studies. For Rosi treatment and the diet switch studies, after 13 weeks of HFD, diet was switched to Rosi-containing HFD (60mg/kg diet) or NCD for 3 weeks. HFD group remained on HFD for the 3 weeks. Monocyte tracking experiments were performed as described previously ^74^. Briefly, PKH67-labeled WT monocytes (0.5×10^6^) were adoptively transferred into each HFD WT recipient mouse via intravenous injection. 3 days or 3 weeks after, the presence of PKH67-labeled cells in adipose tissue of recipient mice were assessed by flow cytometry.

### ATM isolation and flow cytometry analysis

ATM preparation and flow cytometry analysis/sorting were performed as described previously ^74^ with minor modifications. Briefly, after measuring the fat mass, adipose tissue was minced with scissors and digested by incubating in collagenase buffer solution (1x HBSS, 10 mM HEPES, 1.5 mg/mL Collagenase IV, and 8.25 μg/mL DNAse I) with shaking for 20-60 min (depending on fat weight) at 37 °C. Digestion was stopped by adding FACS Buffer (1xDPBS, 0.5 % BSA, 5 mM EDTA), and the suspension was filtered through cotton gauze. Erythrocytes were removed by using Red Blood Cell Lysis Buffer. Later, cells were washed and filtered (Corning, Cat# 352235) through a 35μm mesh for FACS staining. To reduce unspecific binding, the Fc receptor was blocked with CD16/32 prior to incubation with primary antibodies for 30 min ∼ 1 h on ice. All mAbs used for flow cytometry were listed in **Table S1**. For intracellular staining (e.g., Ki67), after surface marker and live/dead cell staining, cells were fixed and permeabilized using the Foxp3 Staining Kit according to the manufacturer’s instructions. Live (DAPI^-^ or Live/Dead Aqua), CD45^+^, CD11b^+^, F4/80^+^ ATMs were sorted or analyzed using SONY MA900 Cell Sorter (Sony Biotechnology, San Jose, USA) and FlowJo software 10.6.2 (BD). To collect ATM-CM cells were incubated for 24h in DMEM supplemented by 10% exosome-free FBS.

### Isolation of non-parenchymal liver cell (NPC)

To isolate liver NPCs, mice were anesthetized and perfused with SC-1 buffer (8 g/L NaCl, 0.44 g/L KCl, 88.17 mg/L NaH_2_PO_4_*H_2_0, 120.45 mg/L Na_2_HPO_4_, 2.38 g/L HEPES, 0.35 g/L NaHCO_3_, 1.9 g/L EGTA, 9 g/L Glucose, pH 7.3-7.4) through inferior vena cava (5 mL/min) for 4 min. When liver color is changed to white or beige, perfusion buffer was switch to SC-2 (8 g/L NaCl, 0.44 g/L KCl, 88.17 mg/L NaH_2_PO_4_*H_2_0, 120.45 mg/L Na_2_HPO_4_, 2.38 g/L HEPES, 0.35 g/L NaHCO_3_, 84.3 g/L CaCl_2_, pH 7.3-7.4) containing 0.5 mg / mL Collagenase D (Roche, Cat# 11088882001) for 5 min (5 mL/min). Liver was excised from mouse, collected into a 50 mL tube containing 20 mL SC-2, and stirred gently with a scrapper (Corning, Cat# 3008) on ice. Digested liver solution was filtered through a 100 μm mesh (Corning, Cat# 352360) and centrifuged at 100 g for 5 min at 4°C to separate hepatocytes (pellets). Supernatant containing NPCs was centrifuged at 1,500 g for 10 min at 4°C to pellet NPCs. To remove stellate cells, the pellet was resuspended with 30 % Histodenz (Sigma, Cat# D2158) and spun down at 2,000 g for 20 min without break at RT. NPCs were further enriched with a 30 % percoll and pelleted by spinning 10 min at 600 g at RT. Pellet was resuspended in FACS Buffer and used for further antibody staining and flow cytometry analysis.

### Isolation and characterization of sEVs

If frozen, macrophage CM was slowly thawed and maintained on ice. To remove residual cells or debris, macrophage CM was centrifuged at 2,000 g for 20 min at 4 °C and filtered through a 0.22 μM filter into Ultra-centrifuge tubes (U-tubes: 25 x 89 mm, 32 mL tubes) (Beckman Coulter, Cat# 355631). To eliminate larger extracellular vesicles, balanced U-tubes were first centrifuged at 16,000 g for 40 min, 4°C using the Optima XPN-80 Ultracentrifuge from Beckman Coulter. Supernatant was transferred into new U-tubes, and smaller extracellular vesicles (including exosomes) were spun down with maximal speed at 120,000 g for 4 h at 4°C. sEV pellets were resuspended in PBS and stored at -80 °C until use. Particle concentration and size of sEVs were measured by using NanoSight NS300 (Malvern Instruments). Expression of exosomal marker proteins, such as CD9, CD63, CD81, and Alix, were assessed by Western Blots. For treatment of cells with sEVs, 1×10^8^ particles were added per well in 24 well plates. 24h later, glucose uptake activity was measured.

### L6 myotubes differentiation

L6 GLUT4myc cells (Kerafast, Cat# ESK202-FP) were seeded 2×10^4^ cells per well in 24 well plates and cultured in MEM-α supplemented by 10% FBS, 1% GlutaMax, and antibiotics. At 80% confluency, to induce differentiation to myotubes, medium was changed to MEM-α containing 2% FBS and maintained for >10 days.

### Glucose uptake

After 24h incubation with BMDM-CM or macrophage-derived sEVs, L6 myotubes were starved in serum-free medium (MEM-α supplemented by 0.25 % BSA) for 16h. Cells were washed with Krebs-Ringer Bicarbonate Buffer (HRKB; 137 nM NaCl, 4.8 mM KCl, 1.2 mM KH_2_PO_4_, 1.2 mM MgSO_4_, 2.5 mM CaCl_2_, 0.2% BSA, 16 mM HEPES) and incubated in HRKB. 2-3 h later, cells were stimulated by 200 nM insulin in HKRB for 30 min. ^3^H-2-deoxy-D-glucose (0.1 mM, 0.4 mCi/mL) was added to cells for 10 min. Glucose transport was stopped by adding 15 μM Cytochalasin B (#C6762, Sigma) for 5 min and then washed twice with ice-cold PBS. Cells were then shaken for 30 min in 1 M NaOH to dissolve the cells efficiently. For detecting the protein concentration, an aliquot of the cell solution was collected before neutralization with 1 M HCl. Finally, extracts were transferred to scintillation vials containing scintillation fluid (Ecosint LS-271, national diagnostics). The radioactivity (^3^H) was counted for 2 min per sample, and counts per minute (CPM) were normalized to total protein content. *Ex vivo* glucose uptake activity in primary mouse adipocytes was measured after 24 h serum starvation as described previously ^75^. Briefly, cells were equilibrated in Hepes-Salt buffer (10 mM Hepes, 40 mM potassium chloride, 125 mM sodium chloride, 0.85 mM potassium phosphate monobasic, 1.25 mM sodium phosphate dibasic, 1 mM magnesium chloride, 1 mM calcium chloride, 0.1% fatty acid-free BSA) for 2 h for glucose starvation, treated with or without 100 nM insulin for 20 minutes followed by addition of 2 μCi ^3^H-2-deoxy-D-glucose for 10 minutes at 37 °C. The glucose uptake was halted by washing with ice cold PBS. The cells were lysed with sodium hydroxide, protein concentration measured, samples were neutralized with hydrochloric acid, and samples were counted in scintillation fluid.

### scRNA-seq analysis

Quality control, alignment, and quantification of reads were performed using Cell Ranger v.(5.0.1) software from 10X Genomics. Sequencing reads were aligned to the mouse genome prepared by 10X Genomics (mm10-2020-A). The R package Seurat ^76^ was used for downstream dimensional reduction, clustering, and differential expression analyses. Prior to downstream analyses, cells with a percentage of mitochondrial genes >15% and the number of unique genes per cell <1000 and >6000 were filtered out. After filtering, gene expression levels were log-normalized and scaled using Seurat functions NormalizeData and ScaleData, respectively ^76^. The FindVariableFeatures ^76^ function was used to find the top 2,000 genes by variable dispersion for each condition. Data were integrated using FindIntegrationAnchors ^76^ followed by IntegrateData ^76^ using the first 30 dimensions. Differentially expressed genes were calculated using the Wilcoxon rank sum test implemented by FindMarkers ^76^. ATMs were distinguished from monocytes by higher expression of macrophage markers (such as *Adgre1* and *Fcgr1*) and lower (or no) expression of monocyte markers (*Ly6c2, Plac8, Chil3, Cx3cr1, Ccr2*, and *Ear2* ^47^). RNA velocity was calculated using the run10X function from Velocyto v0.17.17 ^77^ directly on Cell Ranger outputs. Downstream analyses of RNA velocity were handled using R package velocyto.R and Python package velocyto. R Package CytoTRACE v0.3.3 ^78^ was used to predict differentiation states. Pseudotime trajectory analysis was performed using R package monocle3 ^79^. Trajectory plots were calculating using the plot_cells function. Analyses were repeated on various subsets of monocytes, macrophages, and proliferating cells to compare cell type trajectories across conditions.

#### Western Blot Analysis

Cells were homogenized in RIPA buffer supplemented with protease and phosphatase inhibitors. Protein bands were detected with the SuperSignal West Pico (ThermoFisher Scientific, Cat# 34580) or Femto Chemiluminescent Substrate (ThermoFisher Scientific, Cat# 34095) in the ChemiDoc XRS imaging system and Image Lab 6.1 software (BioRad).

#### Quantitative RT-PCR Analysis

RNA was extracted by using the RNA extraction protocol with Trizol (Invitrogen, Cat# 15596026) or RNeasy Plus Universal Mini kit (QIAGEN, Cat# 73404), according to manufacturers’ instructions. Reverse transcription was performed with the High Capacity cDNA Reverse Transcription Kit (ThermoFisher Scientific, Cat# 4368813). Quantitative realtime RT-PCR was conducted using PerfeCTa SYBR Green FastMiX (Quantabio VWR, Cat# 95073-05K) and primers listed in Table S2 in the StepOnePlus Real-Time PCR Systems (Thermo Fisher Scientific).

### BMDM preparation

After removing all muscles from the hind legs of ≥ 8-week-old male C57BL/6J mice, femur and tibia bones were isolated and flushed out with sterile phosphate-buffered saline (PBS) supplemented by 2% fetal bovine serum (FBS) (2%FBS-PBS). Bone marrow cells contained in flushed PBS were filtered through a 40 μm strainer (Falcon, Cat# 352340) and spun down for 5 min at 500g. Cell pellets were incubated in Red Cell Lysis Buffer (154 mM NH_4_Cl, 10 mM KHCO_3_, 0.1 mM EDTA) for 3 min and then filled with 2%FBS-PBS. After a 5 min centrifugation at 500 g, cells were resuspended in complete RPMI supplemented by 10% FBS, GlutaMax (Thermofisher, Cat# 35050061), and antibiotics. To induce macrophage (BMDM) differentiation, 20 ng/mL recombinant mouse M-CSF (Biolegend, Cat# 576406) was added the culture media. Every 2 days, half of the culture medium was removed and the same volume of fresh RPMI containing M-CSF was added.

To collected BMDM-CM, 24h after treatment with CXCL12, cells were washed three times and incubated in RPMI containing 10% exosome-depleted FBS (Gibco, Cat# A2710801). CM was collected after another 24h and stored at - 80°C.

### Monocyte chemotaxis

Monocytes isolated from healthy wild type mouse bone marrow and incubated in serum-free RPMI medium in 5% CO_2_ incubation for 4 h at 37°C. Chemotaxis activity was measured in Transwell Plates (Corning Costar, Cat# No 3422) with 8 mm pore size (24 well plate). In lower chamber, serum-free RPMI medium containing or not containing 10 nM MCP-1 and/or 100 nM CCL26 was added. In upper chamber, 4 ×10^4^ monocytes in 200uL of serum-free RPMI medium was added. 3h later, cells migrated to the lower side of upper chamber and attached to membrane were washed with ice-cold PBS twice and fixed with ice-cold 100% methanol for 10 min. After staining with 0.05% crystal violet solution in 25% methanol for 10 min, the number of migrated monocytes was counted.

### Caspase-3/7 activity

Caspase-3/7 activity was measured according to the manufacturer’s instruction (AAT Bioquest, Cat# 13507). Briefly, 4 x 10^4^ BMDMs in 100 µL of RPMI medium supplemented by 1% GlutaMax, and 10% FBS were seeded in each well of 96-well plate (Corning, Cat# 3603). After 24h incubation in serum free-media, 100 µL caspase-3/7 working solution was added per well and the plate was incubated for 1 hour at room temperature, protected from light. Absorbance at 490 nm was measured using a plate reader.

### TBARS (lipid peroxide) measurement

Intracellualr TBARS levels in ATMs were measured according to the manufacturer’s instruction (Cayman Chemicals, Cat#10009055).

### Data availability

All data generated by the present study are included in this article and the Supplemental data. The scRNA-Seq data have been deposited in the NCBI’s Gene Expression Omnibus (GEO) database (GEO GSE267767).

### Study Approval

All animal procedures were approved by the local Animal Care and Use Committee and performed in accordance with the University of California, San Diego Research Guidelines for the Care and Use of Laboratory Animals.

### Statistics

The results are shown as means +/- SEM. All statistical analysis was performed by two-tailed Student’s *t* test or ANOVA, unless indicated; *p* < 0.05 was considered significant. Statistical methods were not used to predetermine necessary sample size, but sample sizes were chosen based on estimates from pilot experiments and previously published results such that appropriate statistical tests could yield significant results. Statistical analyses used in the data presented are described in all legends. Parametric tests are used that assume normal distribution, which we showed to be the case when data were plotted as frequencies. Experiments were not performed in a blinded fashion.

The authors have declared that no conflict of interest exists.

---

**Table S1.** Detailed Information of The Reagents and Resources

| REAGENT or RESOURCE | SOURCE | IDENTIFIER |
| --- | --- | --- |
| **Antibodies** |  |  |
| Anti-SAPK/JNK | Cell signaling | Cat# 9252S; RRID:AB_2250373 |
| Anti-phospho SAPK/JNK, (Thr183/Tyr185) | Cell signaling | Cat# 9255S; RRID:AB_2307321 |
| Anti-p38 MAPK | Cell signaling | Cat#9212S; RRID:AB_330713 |
| Anti-phospho p38 MAPK | Cell signaling | Cat#9211S; RRID:AB_331641 |
| Anti-Erk | Cell signaling | Cat# 9102; RRID:AB_330744 |
| Anti-phospho Erk | Cell signaling | Cat#9101; RRID:AB_331646 |
| Anti-Akt | Cell signaling | Cat# 4685; RRID: AB_2225340 |
| Anti-phospho Akt (S473) | Cell signaling | Cat# 4060; RRID: AB_2315049 |
| Anti-β-Actin | Sigma | Cat# A2228; RRID:AB_476697 |
| Anti-α-Tubulin | Abcam | Cat# ab4074; RRID:AB_2288001 |
| Anti-CD45 Brilliant Violet 605 | BioLegend | Cat# 103139, RRID: AB_2562341 |
| Anti-CD11b PerCP-Cyanine5.5 | Thermo Fisher | Cat# 45-0112-82, RRID: AB_953558 |
| Anti-CD11b APC-Cyanine7 | BioLegend | Cat# 101226, RRID: AB_830642 |
| Anti-CD11b PE | Thermo Fisher | Cat# 12-0112-82, RRID: AB_2734869 |
| Anti-F4/80 PE-Cyanine7 | Thermo Fisher | Cat# 25-4801-82, RRID: AB_469653 |
| Anti-F4/80 PE | BioLegend | Cat# 123110, RRID: AB_893486 |
| Anti-CD11c APC | Thermo Fisher | Cat# 17-0114-82, RRID: AB_469346 |
| Anti-Ki67 FITC | Thermo Fisher | Cat# 11-5698-82, RRID: AB_11151330 |
| Anti-Ki67 PE | Thermo Fisher | Cat# 12-5698-82, RRID: AB_11150954 |
| Anti-Ly6C PE | Thermo Fisher | Cat# 560592, RRID: AB_1727556 |
| Anti-CD301b PE/Dazzle | Biolegend | Cat# 146816; RRID: AB_2566027 |
| Anti-Trem2 AF-488 | R&D Systems | Cat# FAB17291G; Source ID: SCR_006140 |
| Anti-CD16/CD32 | Biolegend | Cat# 156604; RRID:AB_2783138 |
| Anti-CXCL12 | Biolegend | Cat# W15149B; RRID:AB_2894525 |
| Anti-CCL26 | Bioss | Cat# bs-15513R |
| **Chemicals, Peptides, and Recombinant Proteins** |  |  |
| Palmitic acid | Sigma | Cat# P9767 |
| Insulin (Norvolin R) | Norvo Nordisk | Cat# 0169-1833-11 |
| 50% Dextrose | Hospira | Cat# 0409-6648-02 |
| MCP-1 | Peptotec | Cat# 250-10 |
| CCL26 | Ray Biotech | Cat# 230-00055 |
| CXCL12 | Peptotec | Cat# 250-20A |
| Collagenase IV | Worthington | Cat# C2139 |
| Collagenase from *Clostridium histolyticum* | Sigma | Cat# C1764 |
| 60% high-fat diet | Research Diets | Cat# D12492 |
| Trizol™ Reagent | ThermoFisher | Cat# 15596026 |
| High Capacity cDNA Reverse Transcription Kit | ThermoFisher Scientific | Cat# 64368813 |
| RBC lysis buffer | eBioscience | Cat# 00-4333-57 |
| Recombinant Mouse M-CSF | Biolegend | Cat# 576406 |
| Endotoxin-low Bovine Serum Albumin | Sigma | Cat# A8806 |
| Free acid-free Bovine Serum Albumin | Sigma | Cat# A7030 |
| LIVE/DEAD™ Fixable Cell Stain Kit | ThermoFisher | Cat# L34966 |
| PKH67 Fluorescent Cell Linker Kits | Sigma | Cat# MINI67-1KT |
| **Critical Commercial Assay Kits** |  |  |
| TBARS assay kit | Cayman Chemicals | Cat# 10009055 |
| Caspase-Glo® 3/7 Assay System | AAT Bioquest | Cat# 13507 |
| **Experimental Models: Cell Lines** |  |  |
| L6-GLUT4myc | Kerafast | Cat# ESK202-FP |
| **Software and Algorithms** |  |  |
| GraphPad Prism v.10 | GraphPad Software | GraphPad Prism, RRID: SCR_002798 |
| FlowJo | FlowJo | FlowJo, RRID: SCR_008520 |

**Table S2.**
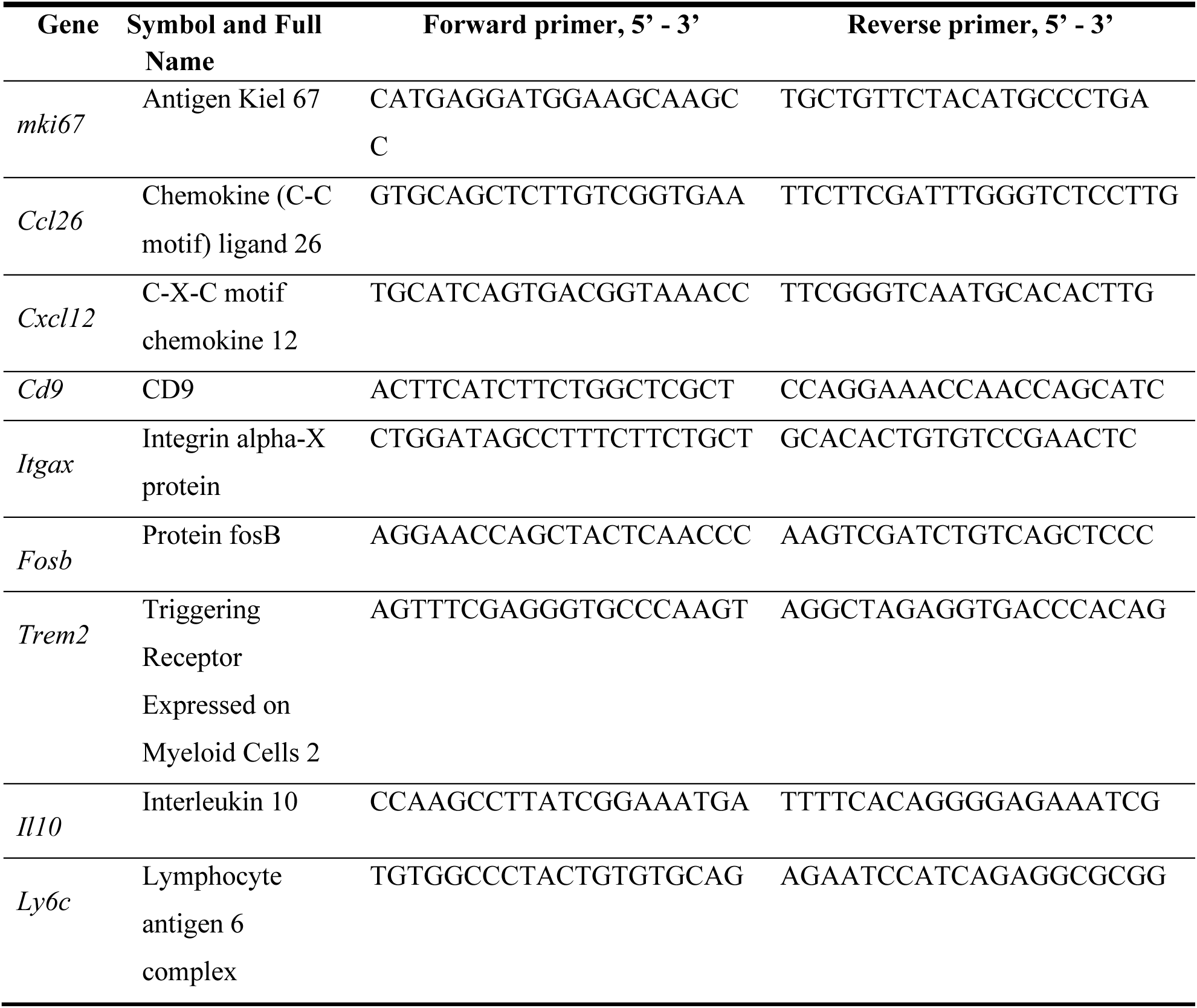
PCR Primer Sequences Gene Symbol and Full

## Supplementary Figure Legends

**Figure S1.**
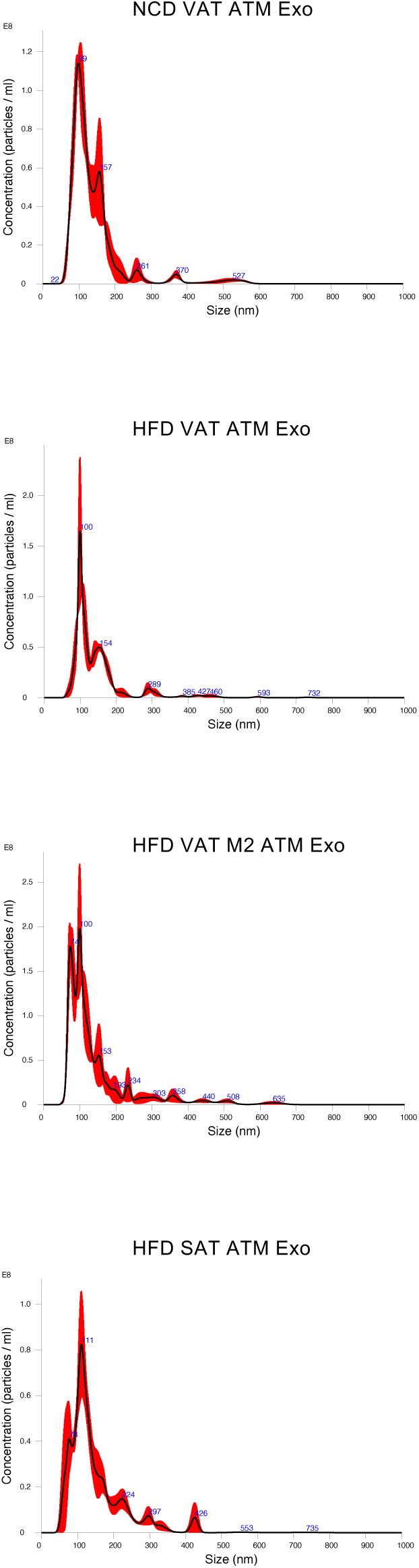
Size distribution of ATM-derived sEVs. To examine whether this effect of ATM sEVs is different in SAT vs VAT, sEVs released from ATMs isolated from subcutaneous inguinal and visceral epididymal white adipose tissue of NCD healthy and HFD obese mice were purified. The sizes and the numbers of ATM-derived sEV particles were validated by by NanoSight. Expression of specific exosomal marker proteins, including Alix, CD63, CD9, and CD81, were tested by Western blots (data not shown).

**Figure S2.**
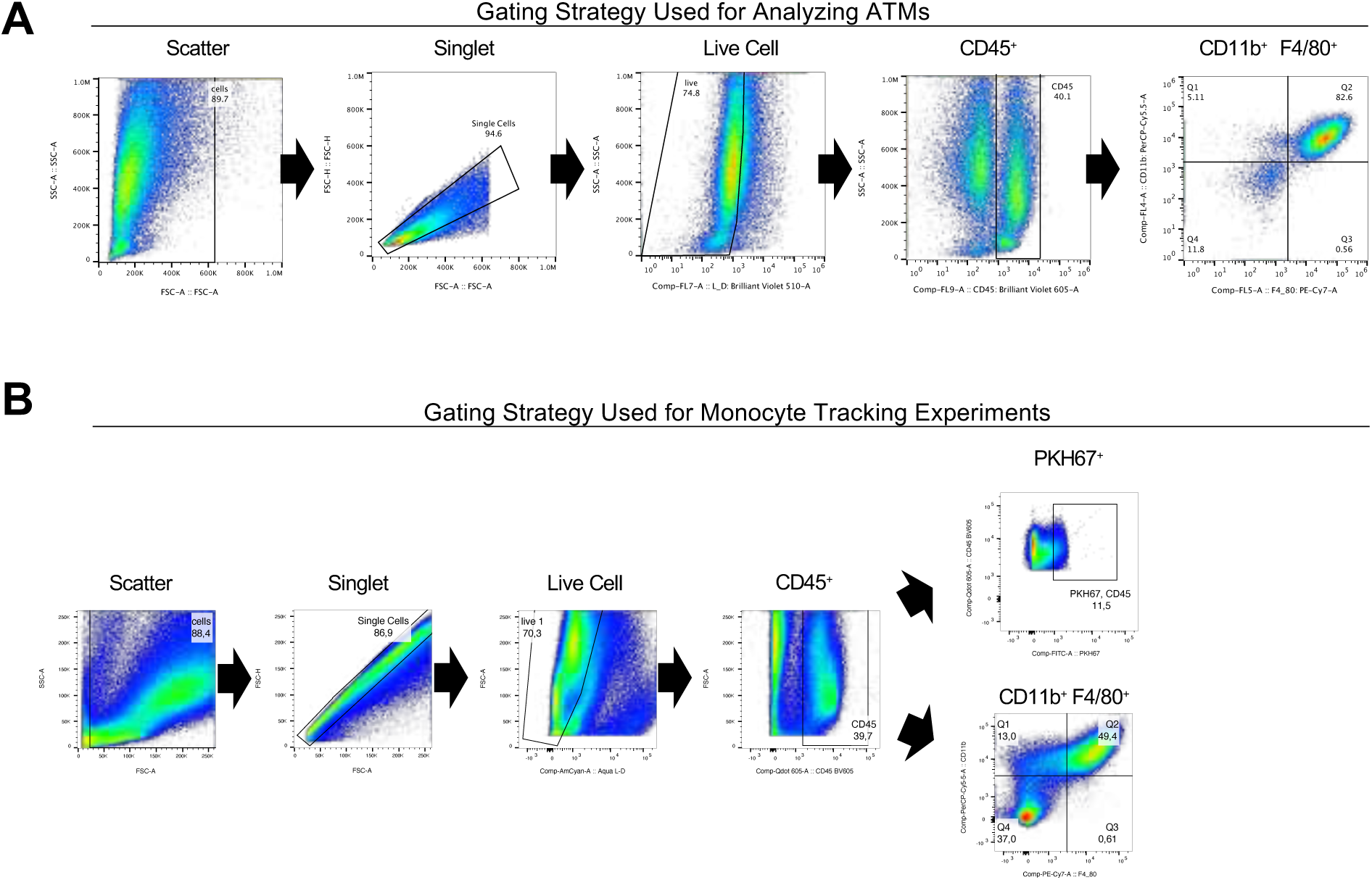
Gating strategy for flow cytometry analysis of adipose tissue monocytes and ATMs in visceral and subcutaneous adipose tissue. (**A**) Gating strategy for flow cytometry data in Figure 1A-1D. (**B**) Gating strategy for flow cytometry data in Figure 1F-1H.

**Figure S3.**
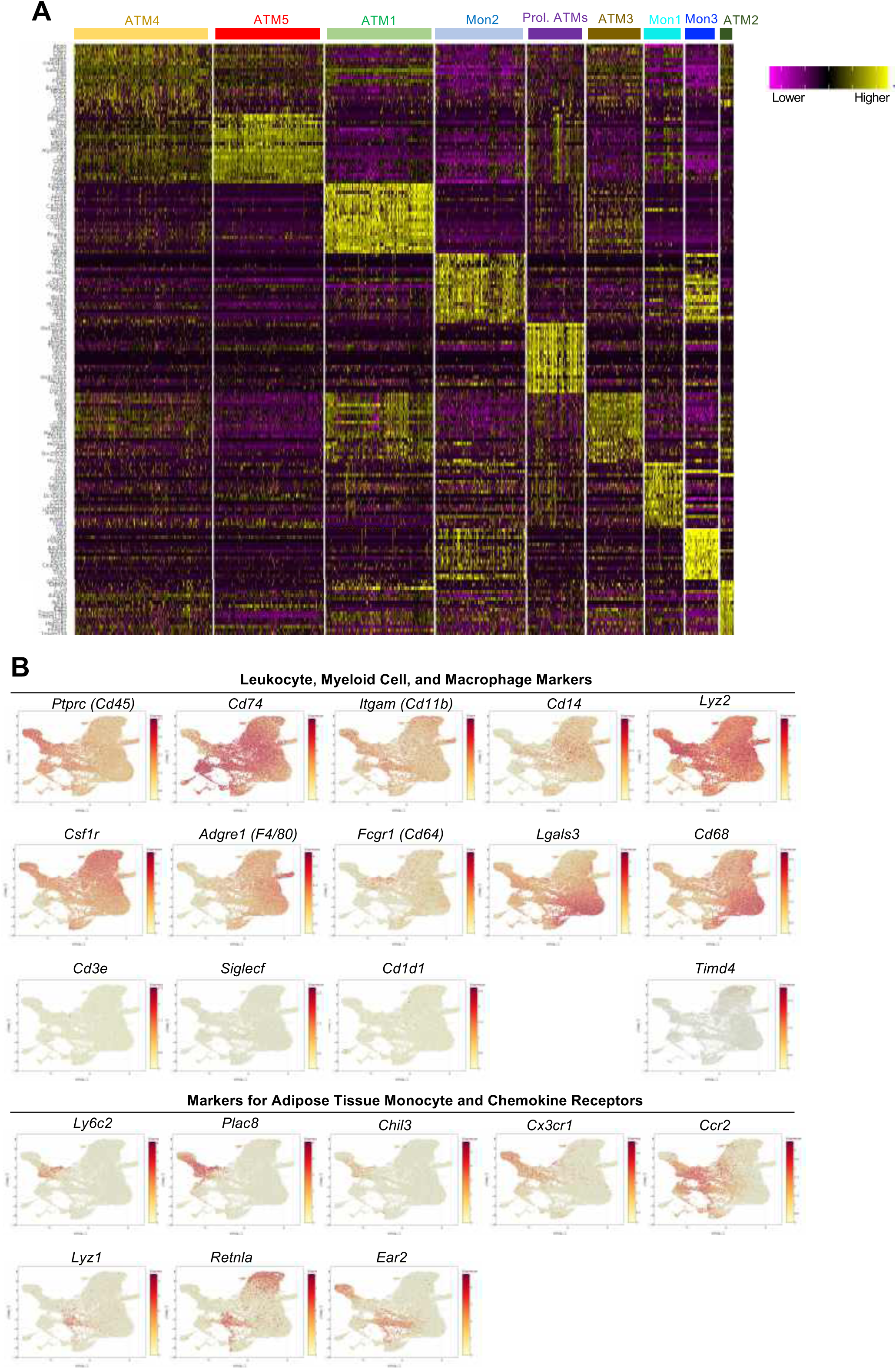
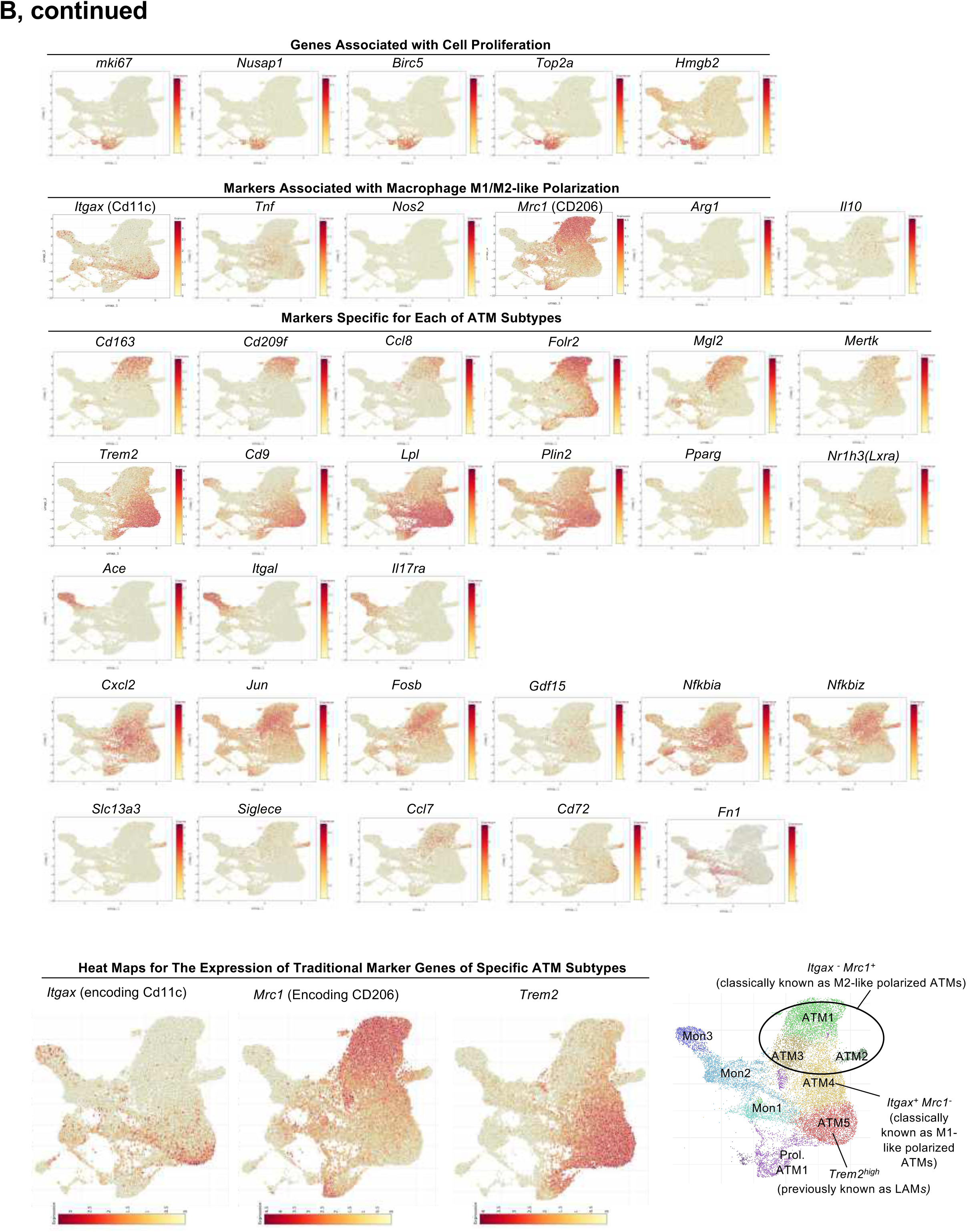
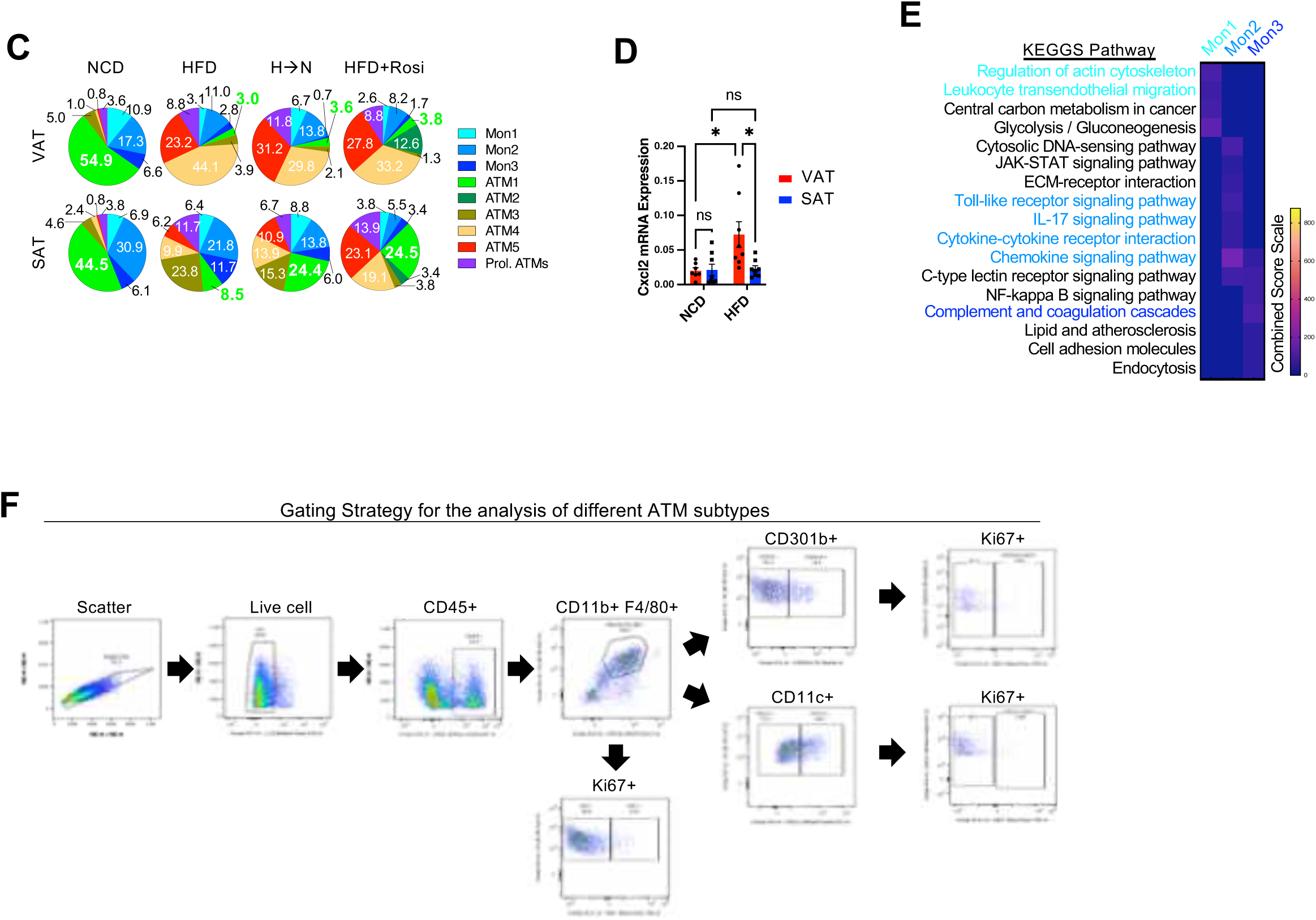
Characterization of ATM and monocyte clusters. (**A**) Heat map showing cluster-specific marker gene expression in different adipose tissue monocyte and ATM clusters. (**B**) Heat map showing cell type or macrophage subtype-specific marker gene expression in different adipose tissue monocyte and ATM clusters. All eight monocyte/macrophage clusters were shared by both SAT and VAT depots and there were no adipose tissue depot-specific clusters across all 4 diet and/or treatment conditions. Despite these commonalities, diet-induced dynamic changes in the proportion of each ATM and monocyte cluster was highly biased by anatomical location of the adipose tissue. The relatively small ATM2 cluster only appeared in Rosi-treated VAT and SAT. (**C**) Proportion of ATM and monocyte clusters in different diet and/or treatment conditions. Inflammatory gene expression was substantially lower in all three monocyte clusters (Mon1, 2, and 3) compared with ATM3 and ATM4. (**D**) *Cxcl2* expression in SAT and VAT of NCD and HFD mice. (**E**) Heatmap showing relative enrichment of pathways in three different adipose tissue monocyte clusters compared with ATMs (scale is the same as the heatmap shown in Figure 3C). (**F**) Gating strategy for flow cytometry analysis of different ATM subtypes (Figure 3I-3K).

**Figure S4.**
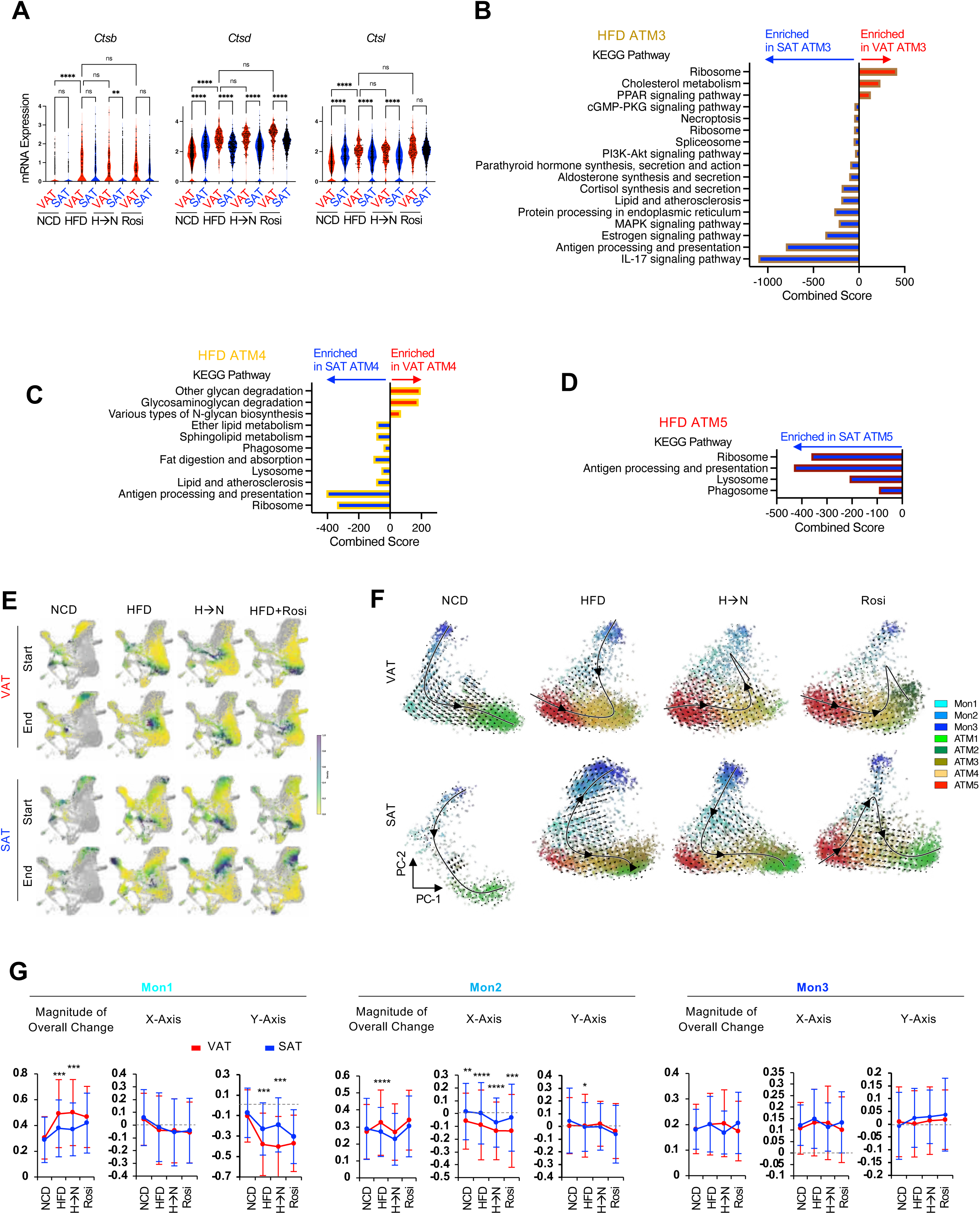
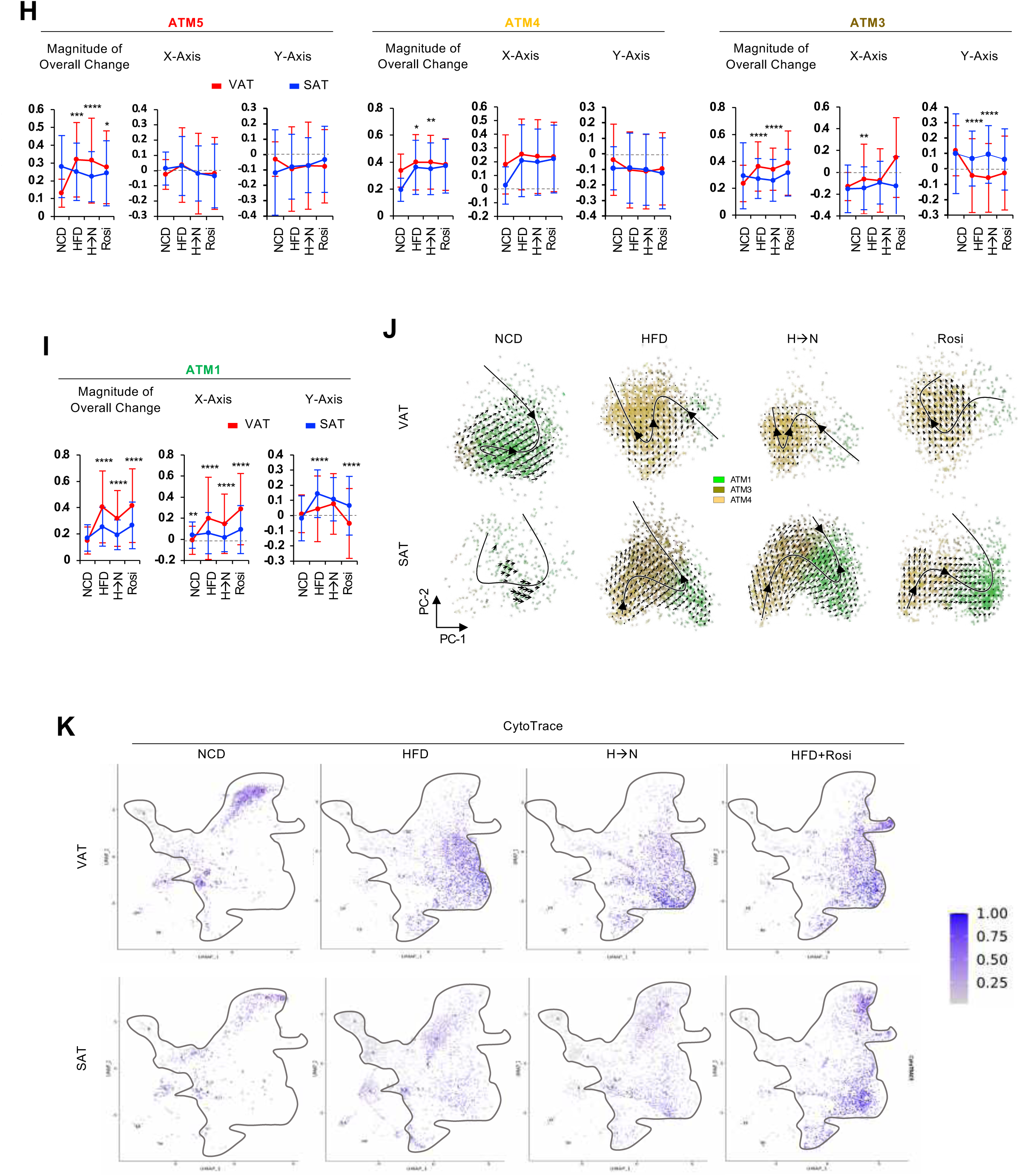
Gene expression and pathways enriched in ATMs from visceral versus subcutaneous adipose tissue. (**A**) Expression of genes involved in lysosomal activity in ATM1 from SAT and VAT of mice fed NCD, HFD, HFD➔NCD switch diet, or HFD mice treated with rosiglitazone (Rosi). (**B-D**) Pathways enriched in visceral or subcutaneous ATM3 (**B**), ATM3 (**C**), and ATM5 (**D**) from HFD mice. (**E-I**) *In silico* cell lineage tracing using RNA Velocity analysis in adipose tissue monocytes and ATMs in mice fed NCD, HFD, the HFD➔NCD switch diet (SwD) or in HFD mice treated with rosiglitazone (Rosi). (**E**) Color-coded UMAP showing the start and end points (Markov transition) of adipose tissue monocyte and ATM differentiation. (**F**) PCA plots showing cell movement trajectory of adipose tissue monocytes and ATMs. Each arrow indicates vectoral changes (size and direction) of transcriptomic signatures in nearest cells. The principle curve (black) indicates integrated directional changes in overall cell movement. Quantitative changes in transcriptomic (vectoral) movement of monocytes (**G**) and ATMs (**H, I**) in panel F were plotted in panel G-I. (**J**) PCA plots showing cell movement trajectory focused on ATM1, ATM3, and ATM4. (**K**) CytoTrace analysis of cell maturity in adipose tissue monocytes and ATMs in SAT and VAT of mice fed HFD, HFD➔NCD switch diet (SwD) or HFD mice treated with rosiglitazone (Rosi). \**P* < 0.05, \*\**P* < 0.01, \*\*\**P* < 0.001, \*\*\*\**P* < 0.0001. ns, not significant. Statistical analysis was performed by 1-way ANOVA with Tukey’s multiple comparison tests. \**P* < 0.05, \*\**P* < 0.01, \*\*\**P* < 0.001, \*\*\*\**P* < 0.0001, SAT vs VAT. Statistical analysis was performed by 1-way ANOVA with Tukey’s multiple comparison tests.

**Figure S5.**
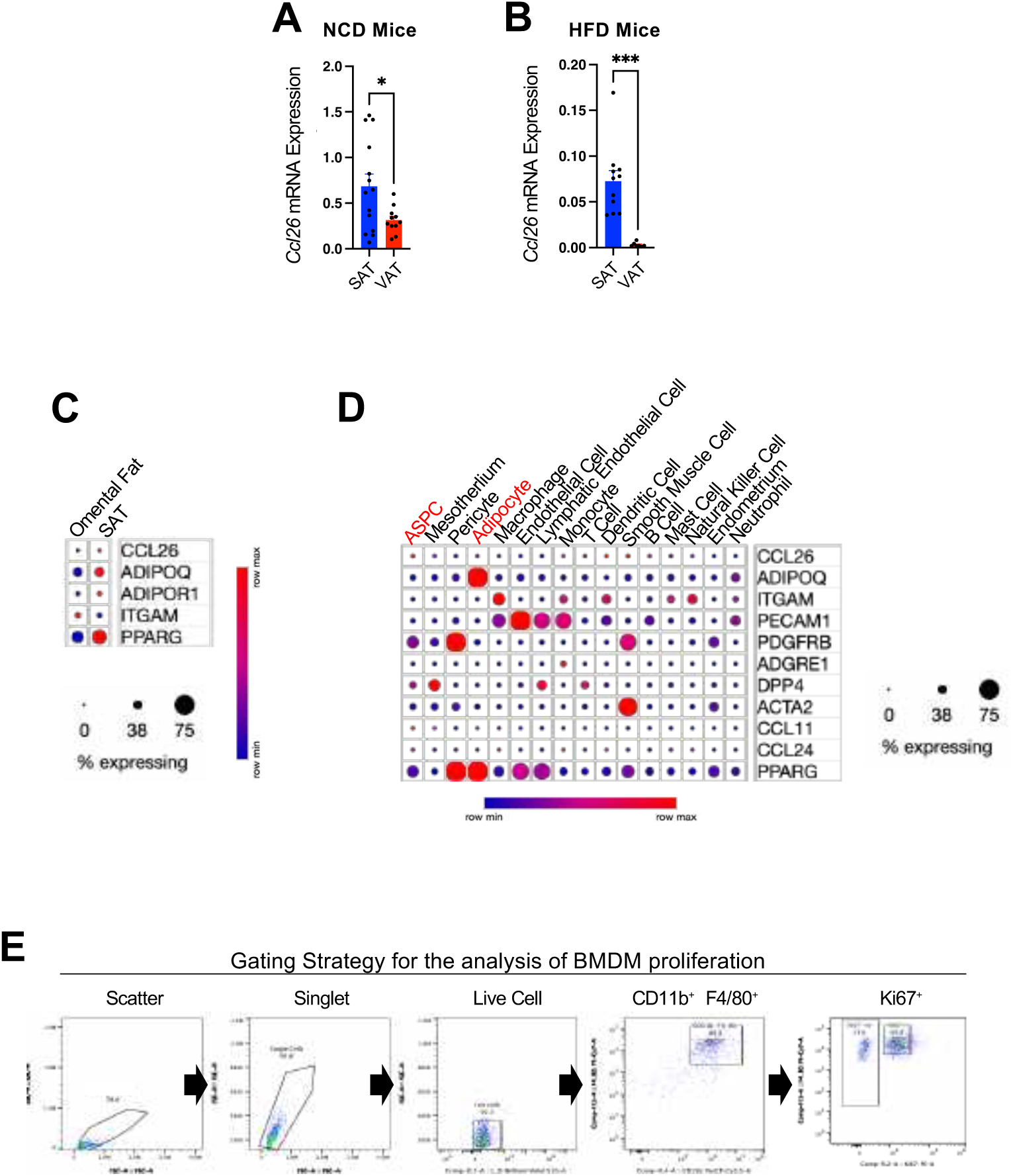
Adipose tissue *Ccl26* expression on NCD and HFD. (**A-B**) *Ccl26* expression in SAT and VAT of NCD (**A**) or HFD (**B**) mice. (**C-D**) snRNA-seq data analysis of *CCL26* expression in human SAT and omental adipose tissue ^50^. Dot plots show relative mRNA expression of *CCL26* in whole human SAT and omental adipose tissue (**C**) and in different cell types consisting adipose tissue (**D**), including adipose stem and progenitor cells (ASPC) and adipocytes. Cell type specific markers for adipocytes (PPARG, ADIPOQ), myeloid lineage cells (ITGAM), endothelial cells (PECAM1), pericytes (PDGFRB), and smooth muscle cells (SMC; ACTA2) as comparators. (**E**) Gating strategy for flow cytometry data in Figure 5I. \**P* < 0.05, \*\*\**P* < 0.001. n.s., not significant. Statistical analysis was performed by the student t-tests.

**Figure S6.**
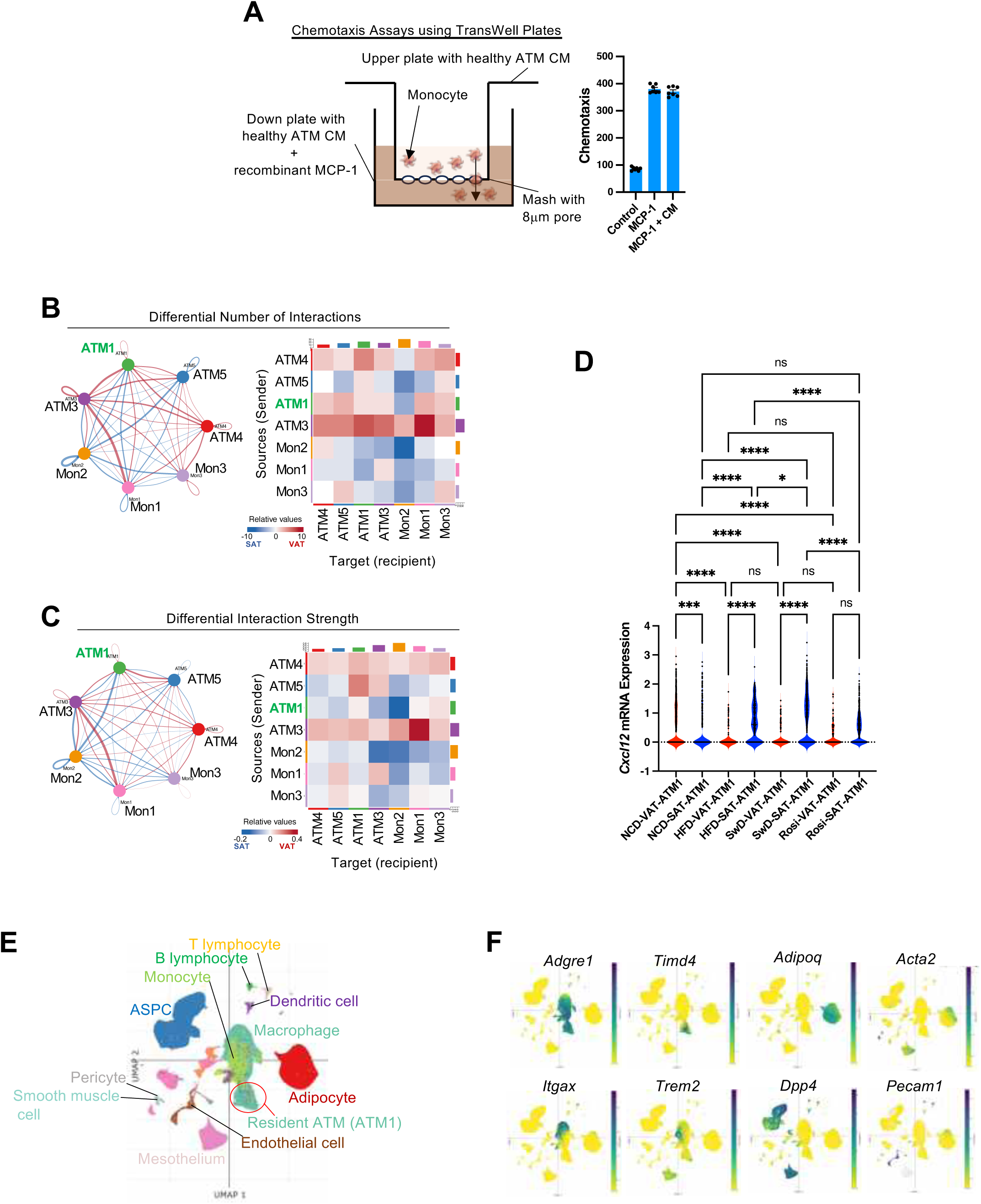
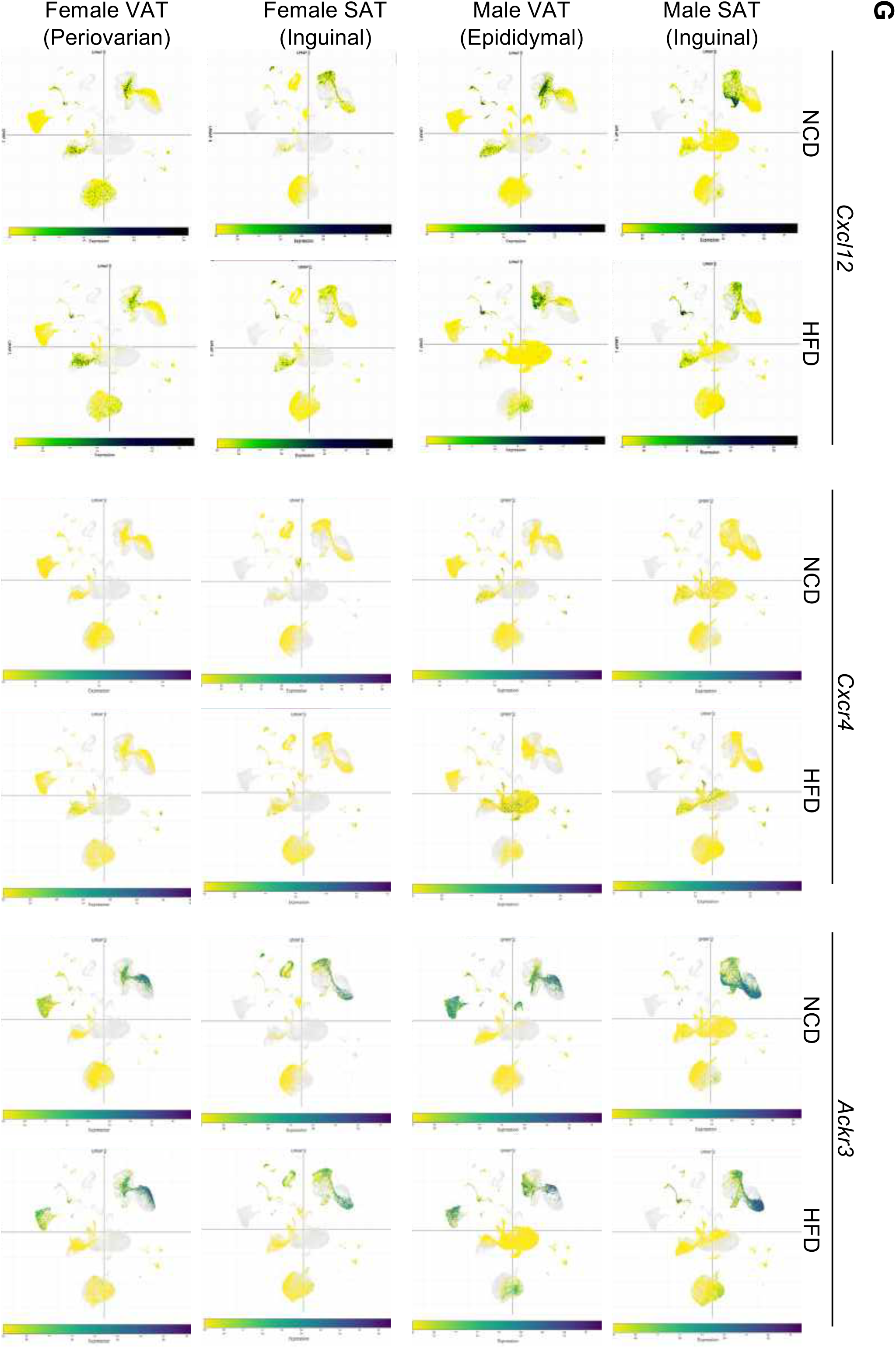
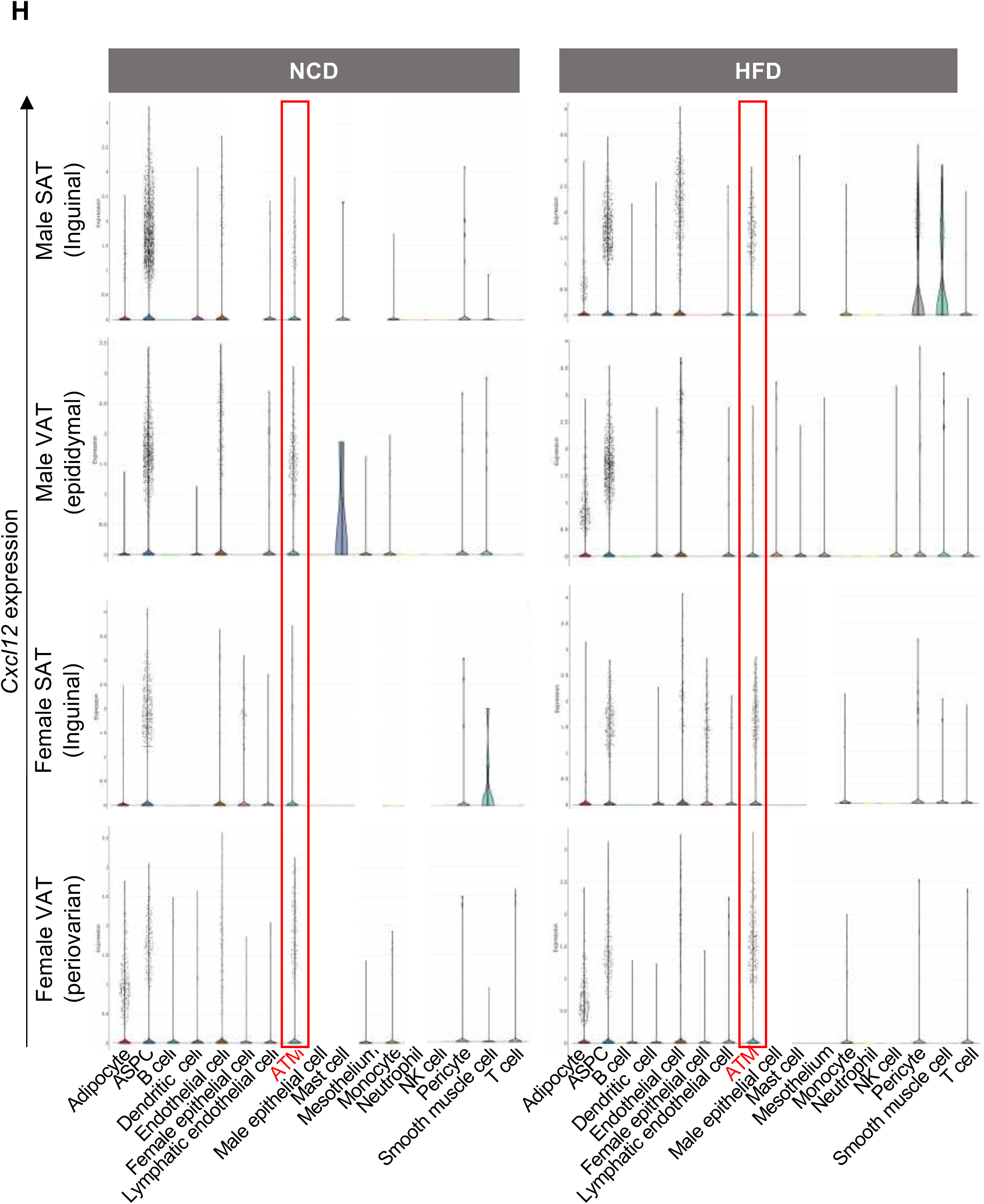
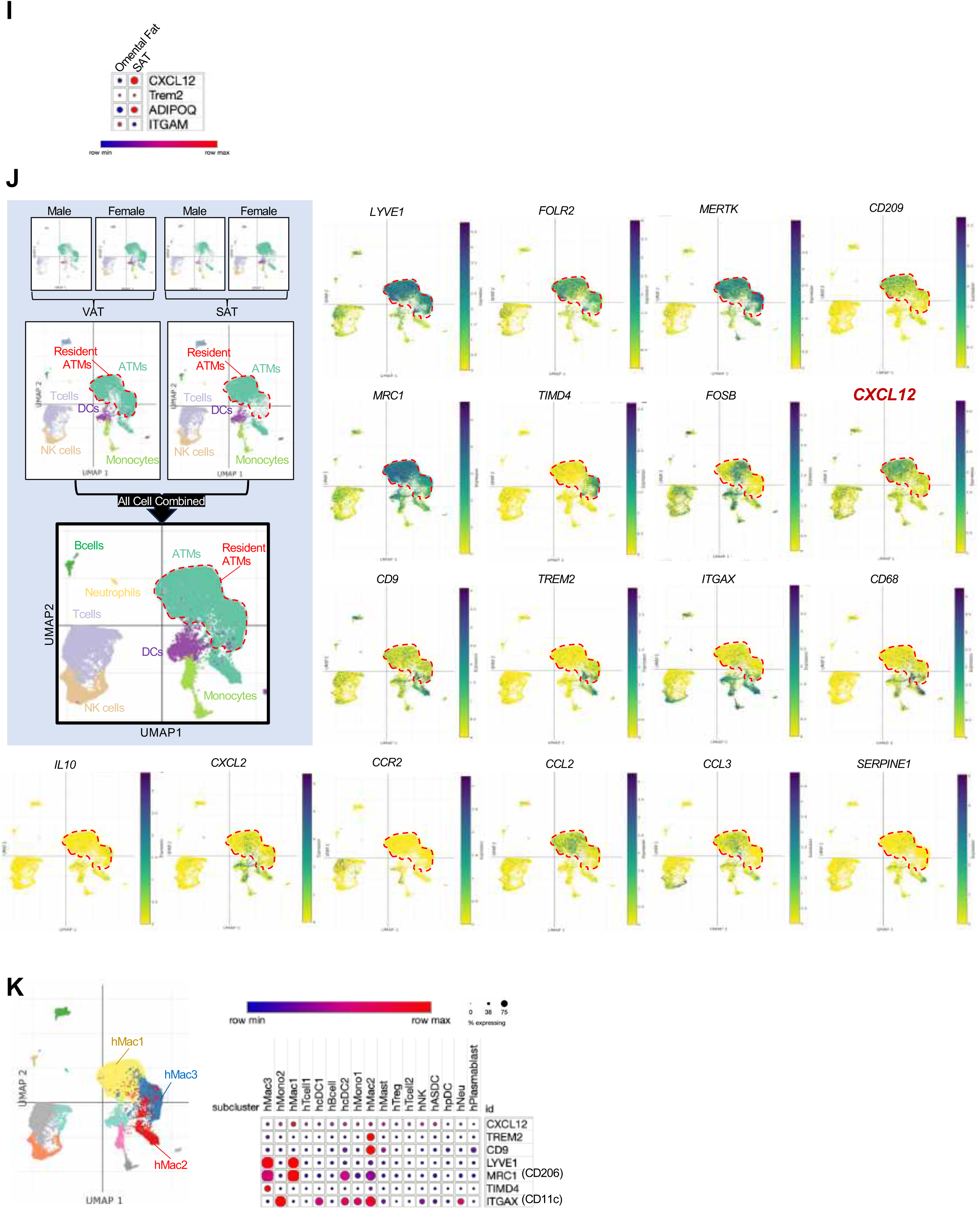
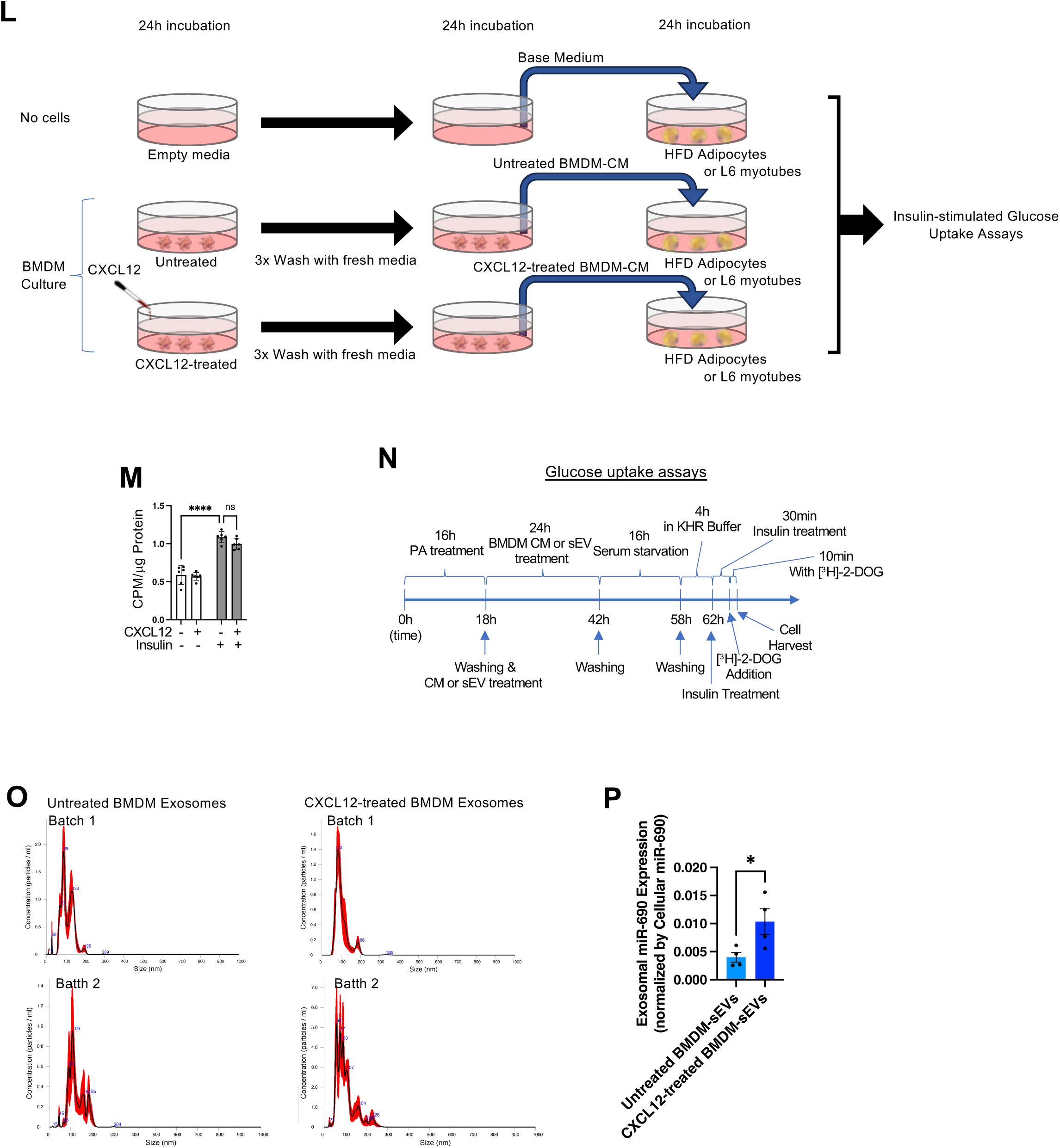
Adipose tissue *Cxcl12* expression and the effect of CXCL12 treatment on BMDM sEV secretion. (**A**) Monocyte chemotaxis assays using MCP-1, +/- healthy ATM CM. As illustrated on left, the effect of healthy ATM-CM on monocyte chemotaxis to MCP-1 was assessed using Tranwell plates. The result was plotted on right. (**B-C**) CellChat analysis of interaction between different adipose tissue monocyte and ATM clusters in HFD mice. Heatmaps show the number (B) and the strength (C) of interaction between individual adipose tissue monocyte and ATM clusters enriched inVAT (red) compared with SAT (blue). (**D**) mRNA expression of *Cxcl12* in ATM1 cells from VAT and SAT of mice fed NCD, HFD, or the switch diet from HFD to NCD, or VAT and SAT of HFD mice treated with Rosi. (**E-H**) Single nuclear RNA-seq analysis of *Cxcl12* and its receptors (*Cxcr4* and *Ackr3*) in SAT and VAT of male and female mice fed NCD or HFD. (**E**) UMAP representation of different cell clusters in all cells from all samples. ASPC, adipose stem cells and progenitor cells. (**F**) Heat map showing the expression of cell type-specific marker genes. (**G**) Heat maps showing the expression of *Cxcl12* and its receptors (*Cxcr4* and *Ackr3*) in SAT and VAT of male and female mice fed NCD or HFD. (**H**) Violin plots showing the expression of *Cxcl12* in different cell types from SAT and VAT of male and female mice fed NCD or HFD. (**I-K**) snRNA-seq data analysis of *CXCL12* expression in ATMs from human adipose tissue ^50^. (**I**) Dot plot showing relative mRNA expression of *CXCL12* in human SAT and VAT. (**J**) *CXCL12* expression in all immune cells. UMAP representation of all immune cells in human VAT and SAT is shown in left upper panel (in boxed area). Heatmaps showing the expression of *CXCL12*, resident (*LYVE1, FOLR2, MERTK, CD209, MRC1, TIMD4*) and recruited (*CD9, TREM2, ITGAX, CCR2*) macrophage marker genes, and pro-inflammatory (*CCL2, CCL3, SERPINE1*) and anti-inflammatory (*IL10*) cytokines are presented on the right. (**K**) A dot plot showing relative expression of *CXCL12* in each of immune cell types, including three different ATM subclusters [hMac1 (*FOSB^+^ LYVE1^+^ TREM2*^-^ *ITGAX*^-^ resident ATMs), hMac2 (*TIMD4^-^ LYVE1^-^ TREM2*^+^ *CD9*^+^ *ITGAX*^+^ recruited ATMs), and hMac3 (*TIMD4^+^ LYVE1^+^ TREM2*^-^ *ITGAX*^-^ resident ATMs)] (right). Color-coded UMAP showing three different ATM clusters (hMac1, hMac2, and hMac3) is shown on the left. (**L**) Schematic representation of experiments in Figure 6K to 6O. (**M**) Glucose uptake by primary adipocytes from HFD/obese WT mice. Before measurements, adipocytes were treated with or without 10 ng/ml CXCL12 for 24 (n=6 wells/group). (**N**) Schematic representation of glucose-uptake assays in Figure 6L-6N. (**O**) NanoSight analyses of sEV particle size of BMDM-derived sEVs used in Figure 6N-6S. (**P**) Relative miR-690 expression in sEVs, normalized by intracellular miR-690 expression in BMDMs treated with or without CXCL12 (n=4 wells/group). \**P* < 0.05, \*\*\**P* < 0.001, \*\*\*\**P* < 0.0001. Statistical analysis was performed by 1-way ANOVA with Tukey’s multiple comparison tests.

